# A hydrophobic funnel governs monovalent cation selectivity in the ion channel TRPM5

**DOI:** 10.1101/2023.01.11.523625

**Authors:** Callum M. Ives, Alp Tegin Şahin, Neil J. Thomson, Ulrich Zachariae

## Abstract

A key capability of ion channels is the facilitation of selective permeation of certain ionic species across cellular membranes at high rates. Due to their physiological significance, ion channels are of great pharmaceutical interest as drug targets. The polymodal signal-detecting Transient Receptor Potential (TRP) superfamily of ion channels form a particularly promising group of drug targets. While most members of this family permeate a broad range of cations including Ca^2+^, TRPM4 and TRPM5 are unique due to their strong monovalent-selectivity and impermeability for divalent cations. Here, we investigated the mechanistic basis for their unique monovalent-selectivity by *in silico* electrophysiology simulations of TRPM5. Our simulations reveal an unusual mechanism of cation selectivity, which is underpinned by the function of the central channel cavity rather than the selectivity filter. Our results suggest that a subtle hydrophobic barrier at the cavity entrance ("hydrophobic funnel") enables monovalent, but not divalent cations to pass and occupy the cavity at physiologically relevant membrane voltages. Monovalent cations then permeate efficiently by a co-operative, distant knock-on mechanism between two binding regions in the extracellular pore vestibule and the central cavity. By contrast, divalent cations do not enter or interact favourably with the channel cavity due to its raised hydrophobicity. Hydrophilic mutations in the transition zone between the selectivity filter and the central channel cavity abolish the barrier for divalent cations, enabling both monovalent and divalent cations to traverse TRPM5.

## Introduction

The translocation of ions across cellular and organellar membranes via ion channels is essential to ensure cellular ionic homeostasis and provides a key pathway of intra- and intercellular communication. Ion channels catalyse the permeation of ions across the membrane up to an order of 10^8^ ions per second, while at the same time often displaying strict selectivity for particular ionic species (***Dudev and Lim, 2014***; ***Roux, 2017***; ***Kopec et al., 2018***). The transient receptor potential (TRP) super-family of ion channels comprises a large group of cation-selective channels that are implicated in a wide range of physiological processes (***Ramsey et al., 2006***; ***Khalil et al., 2018***). Due to their physiological importance, TRP channels are associated with a large number of pathological conditions (***Nilius, 2007***), including in the aetiology of several rare, genetic conditions. Many members of the TRP channel superfamily constitute major pharmaceutical target proteins (***Moran, 2018***; ***Koivisto et al., 2022***).

Within the TRP channel superfamily, TRPM (transient receptor potential melastatin) channels form the largest subfamily, consisting of eight members (TRPM1-8) (***Nilius and Owsianik, 2011***; ***Samanta et al., 2018***). TRPM channels assemble as homotetramers, in which each subunit provides six transmembrane helices (S1-S6), a cytosolic *N*-terminus domain composed of four melastatin homology regions, and a cytosolic *C*-terminus coiled-coil domain (***Hilton et al., 2019***; ***van Goor et al., 2020***). In keeping with most members of the TRP superfamily, TRPM channels are described as being cation non-selective, that is, they conduct cations but do not differentiate substantially between cationic species. However in the TRPM subfamily, TRPM4 and TRPM5 are exceptions to this observation, since both channels are selective for monovalent cations and impermeable to divalent cations (***Owsianik et al., 2006***). Thereby, TRPM4 and TRPM5 are the only members of the wider TRP superfamily to display selectivity for monovalent cations.

Although TRPM4 and TRPM5 are close homologs, sharing both a high degree of sequence homology and similar biophysical characteristics, there are some variations in their activation mechanisms. For example, while both channels are activated by raised intracellular Ca^2+^ concentrations, TRPM5 is approximately 20-fold more sensitive to Ca^2+^ than TRPM4 (***Ullrich et al., 2005***). Ion conduction through TRPM5 has been implicated in the sensation of sweet, bitter, and umami tastes in type II taste bud cells (***Pérez et al., 2002***; ***Zhang et al., 2003***), and in the secretion of insulin by pancreatic 𝛽-cells (***Brixel et al., 2010***; ***Colsoul et al., 2010***). Consequently, TRPM5 is a potential drug target for a number of conditions, including metabolic conditions such as type II diabetes mellitus (***Vennekens et al., 2018***). Several molecular structures of the TRPM4 and TRPM5 channels have been published to date, however an open-state structure has only been solved for TRPM5 (***Ruan et al., 2021***).

In the present work, we set out to characterise the cation permeation mechanism of the TRPM5 channel, focusing in particular on the basis for its monovalent cation selectivity, by conducting atomistic molecular dynamics (MD) simulations and *in silico* electrophysiology of the open-state structure of *Danio rerio* TRPM5 (***Ruan et al., 2021***) (PDB ID: 7MBS) in solutions of Na^+^, K^+^, and Ca^2+^ ions. We recorded more than 700 individual ion permeation events from over 20 𝜇s of aggregated time from our *in silico* electrophysiology simulations.

Our findings reveal a new mechanism of ion selectivity, based on a hydrophobic barrier at the entrance to the central channel cavity, which shields the cavity from an influx of divalent cations. In this way, the central cavity forms a binding site for monovalent cations, but not for divalent cations. The conduction of monovalent ions thus becomes a synergistic process incorporating cooperativity between multiple binding sites.

## Methods & Materials

### TRPM5 system construction

A truncated TRPM5 simulation system consisting of the membrane-domain of the channel was constructed by using residues 698-1020, including the resolved *N*-acetyl-𝛽-D-glucosamine of the glycosylated N921 residue, of the *Danio rerio* TRPM5 structure (***Ruan et al., 2021***) (PDB ID: 7MBS). We also modelled the bound Ca^2+^ cations occupying the Ca_TMD_ binding sites at E768 and D797 in each subunit, which have been proposed to be implicated in Ca^2+^-dependent activation of TRPM5. The system was built using the CHARMM-GUI server (***Jo et al., 2008***). The charged *N*- and *C*-terminal residues were neutralised by capping with acetyl (ACE) and *N*-methylamide (CT3) groups, respectively. All missing non-terminal residues were modelled using CHARMM-GUI (***Jo et al., 2014***).

The structure was aligned in the membrane using the PPM server (***Lomize et al., 2012***), inserted into a 1-palmitoyl-2-oleoyl-sn-glycerol-3-phosphocholine (POPC) bilayer of 160 x 160 Å size with the CHARMM-GUI membrane builder (***Jo et al., 2007***; ***Wu et al., 2014***), and then solvated. Ions were added with GROMACS 2020.2 (***Abraham et al., 2015***; ***Lindahl et al., 2020***) to neutralise any system charges and add ions to a concentration of either 150 mM NaCl, 150 mM KCl, 150 mM CaCl_2_ (referred to as mono-cationic solutions), or a mixture of 75 mM NaCl and 75 mM CaCl_2_ (referred to as di-cationic solutions). In the case of simulations containing Ca^2+^, the standard CHARMM36m parameters for Ca^2+^ ions were then replaced with the multi-site Ca^2+^ of Zhang *et al*. (***Zhang et al., 2020***). This multi-site model has used been used to investigate Ca^2+^ permeation in a number of channels, including including the type-1 ryanodine receptor ***Zhang et al.*** (***2020***); ***Liu et al.*** (***2021***), AMPA receptors (***Schackert et al., 2022***), the E protein of SARS-CoV-2 (***Antonides et al., 2022***), and TRPV channels (***Liu and Song, 2022***; ***Ives et al., 2023***).

### Molecular dynamics simulation details

All simulations were performed using GROMACS 2020.2 (***Abraham et al., 2015***; ***Lindahl et al., 2020***) or GROMACS 2022 (***Bauer et al., 2022***), together with the CHARMM36m force field for the protein, lipids, and ions (except for Ca^2+^ as noted above) (***Huang et al., 2016***). The TIP3P water model was used to model solvent molecules (***Jorgensen et al., 1983***). The system was minimised and equilibrated using the suggested equilibration input scripts from CHARMM-GUI (***Lee et al., 2016***). In brief, the system was equilibrated using the NPT ensemble for a total time of 1.85 ns with the force constraints on the system components being gradually released over six equilibration steps. The systems were then further equilibrated by performing a 15 ns simulation with no electric field applied. To prevent closing of the lower hydrophobic gate of the pore, harmonic restraints were applied to maintain the distance between the 𝛼-carbon atoms of the lower gate residue I966 of each respective chain. The temperature was maintained at T = 310 K using the Nosé-Hoover thermostat (***Evans and Holian, 1985***), and the pressure was maintained semi-isotropically at 1 bar using the Parrinello-Rahman barostat (***Parrinello and Rahman, 1981***). Periodic boundary conditions were used throughout the simulations. Long-range electrostatic interactions were modelled using the particle-mesh Ewald method (***Darden et al., 1993***) with a cut-off of 12 Å. The LINCS algorithm (***Hess et al., 1997***) was used to constrain bond lengths involving bonds with hydrogen atoms. Hydrogen mass re-partitioning (HMR) of the system was used to allow the use of 4 fs integration time steps in simulations of NaCl solutions. The multi-site Ca^2+^ model used for simulations of CaCl_2_ however is incompatible with a 4-fs time step, and therefore any simulations involving Ca^2+^ cations were performed with HMR but at a time step of 2-fs. A summary of all simulations performed is presented in Table 1, and in more detail in Tables S1 and S2.

**Table 1.**
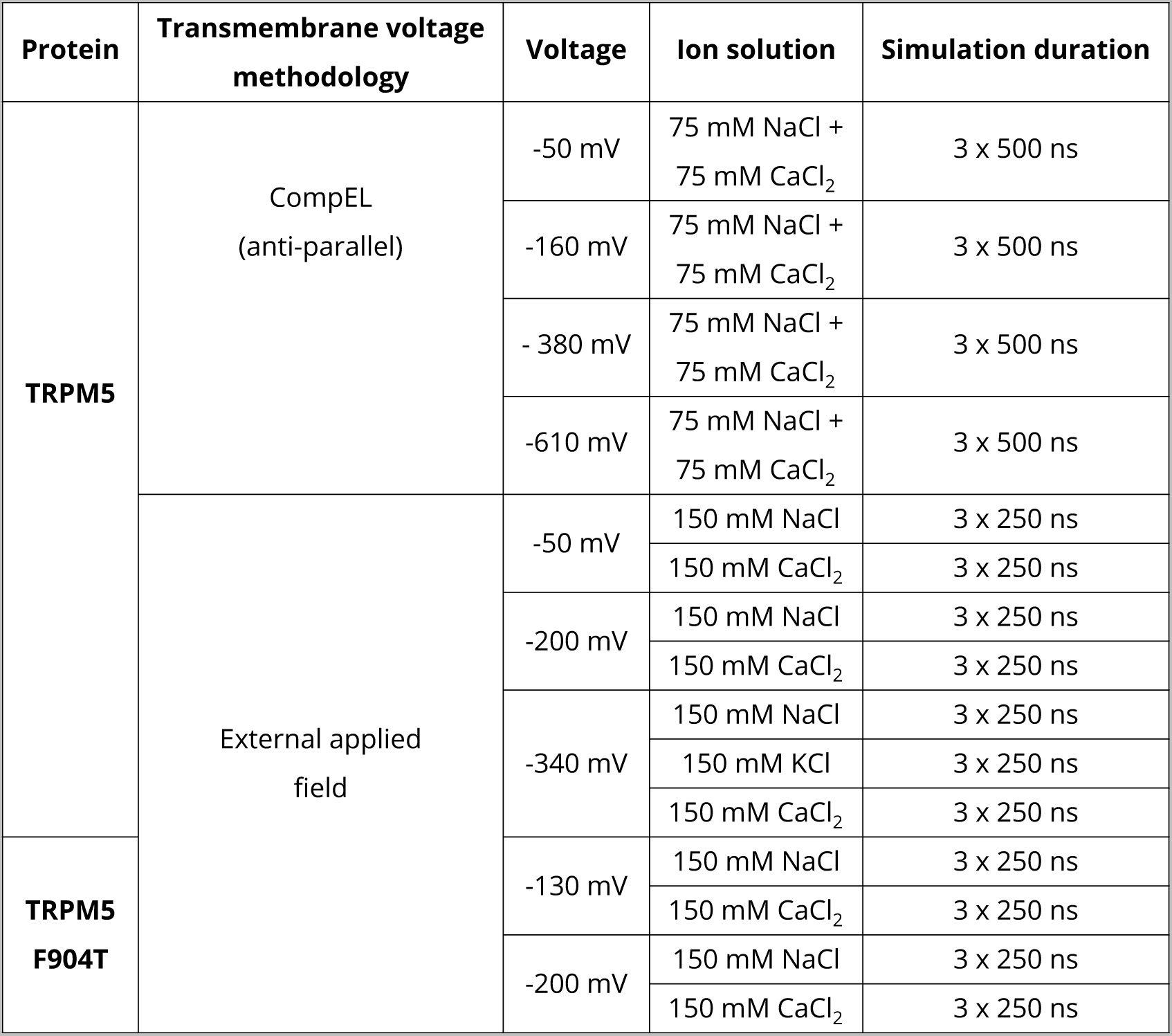
Summary of simulations performed in this study.

### CompEL simulations

We employed the computational electrophysiology (CompEL) protocol (***Kutzner et al., 2011***, ***2016***) of GROMACS to create a transmembrane voltage and drive ion permeation in an anti-parallel double membrane system, such that both channels experienced the same voltage polarity with negative polarity in the intracellular region. Simulations were performed in a di-cationic solution of 75 mM NaCl and 75 mM CaCl_2_ with a range of ionic imbalances (Δq), resulting in membrane voltages of ∼ -50 mV, -130 mV, -380 mV, and -610 mV. To further drive cation permeation, we also generated a neutral ion concentration gradient of 9:1 between the extracellular and intracellular solutions (Figure 1). All CompEL simulations were 500 ns long and repeated three times for each system, resulting in an aggregated simulation time of 3 µs per membrane voltage due to the double channel nature of these simulations.

**Figure 1.**
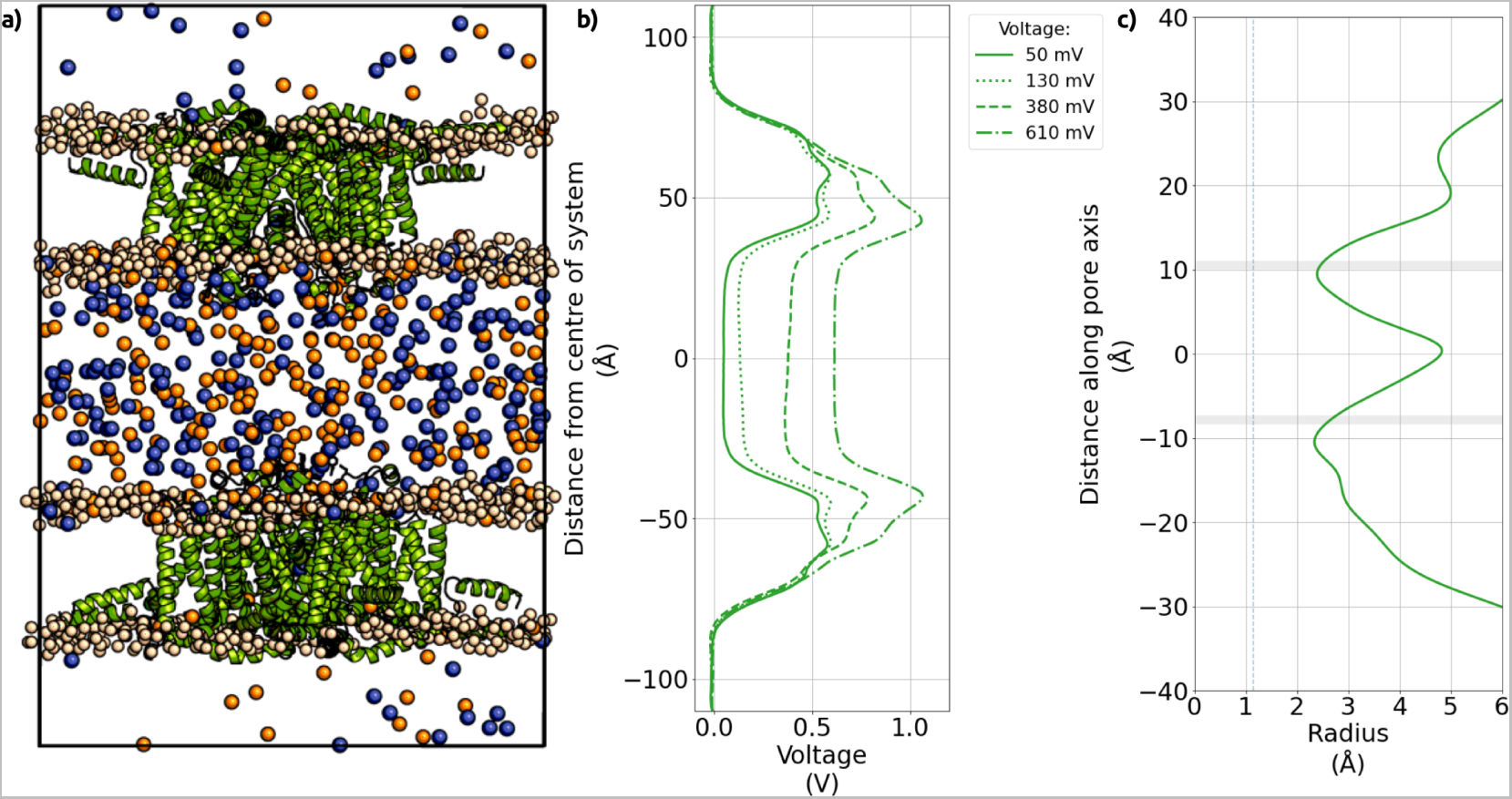
Structure and membrane voltage of CompEL simulations of TRPM5. a) Snapshot of the CompEL system showing the TRPM5 pore domain of *Danio rerio* used in this study inserted into a double bilayer simulation system in an anti-parallel fashion so that both proteins experience identical voltage polarity. Cations within the aqueous compartments are shown as spheres (orange: calcium; blue: sodium), highlighting the 9:1 ion concentration gradient between the compartments. b) The CompEL charge differences we applied across the aqueous compartments (Δq) resulted in transmembrane voltages of ∼ -50 mV, -130 mV, -380 mV, and -610 mV in addition to the concentration gradient. c) Average pore radius of TRPM5 along the pore axis from MD simulations. The regions in grey shade represent the average positions of the major pore constrictions in TRPM5, formed by Q906 (*upper* gate) and I966 (*lower* gate). The dashed line indicates the radius of a completely dehydrated Ca^2+^ ion for comparison.

### External applied field simulations

In addition to CompEL simulations, we also performed simulations in mono-cationic solutions of 150 mM NaCl, 150 mM KCl, and 150 mM CaCl_2_, using an applied electric field to produce membrane voltage ***Aksimentiev and Schulten*** (***2005***). Fields of -0.03, -0.0175, or -0.0044 V nm^-1^ were applied, resulting in transmembrane voltage of ∼-340 mV, 2-00 mV, or -50 mV, respectively, with negative polarity in the intracellular region. All applied field simulations were 250 ns long and repeated three times for each system.

### Simulation analysis

Analysis of MD trajectory data was performed using in-house written Python scripts, utilising GROMACS modules (***Abraham et al., 2015***; ***Lindahl et al., 2020***), the SciPy library of tools (***Oliphant, 2007***; ***Pérez and Granger, 2007***; ***Millman and Aivazis, 2011***; ***Van Der Walt et al., 2011***), and MDAnalysis (***Michaud-Agrawal et al., 2011***; ***Gowers et al., 2016***). Analysis of the pore architecture was performed using CHAP (***Rao et al., 2019***). All plots were generated in Python using Matplotlib (***Hunter, 2007***) and Seaborn (***Waskom et al., 2018***). All MD input and analysis scripts used for this study are deposited in a public GitHub repository, available at: https://github.com/cmives/Na_selectivity_mechanism_of_TRPM_channels.

Calculating conductance and selectivity from *in silico* electrophysiology experiments The conductance of the channels (*C_ion_*) was calculated according to Equation 1, where *N_p_*is the number of permeation events, *Q_ion_* is the charge of the permeating ion in Coulomb, *t_traj_* is the length of the trajectory, and *V_tm_* is the transmembrane voltage. The mean conductance and standard error were calculated from overlapping 50 ns windows of the trajectory.

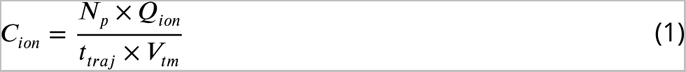

The selectivity (*𝑃_𝑁𝑎_*∕*𝑃_𝐶𝑎_*) from the di-cationic CompEL simulations was calculated as the ratio between the total sum of Na^+^ permeation events and the total sum of Ca^2+^ permeation events across all simulations in a certain voltage or concentration regime.

### Identification of cation binding sites from MD simulations of TRPV channels

Cation binding sites were identified by plotting timeseries of each permeating ion with respect to their position along the pore axis. To further validate these positions, a 3D density mesh was generated for cations within 10 Å of the protein. This analysis was performed on a trajectory of concatenated, three-fold replicated 500 ns simulations in mono-cationic solutions with a voltage of ∼ -50 mV produced by the CompEL method.

### Characterising permeation cooperativity through mutual information with SSI from PENSA

To characterise the level of co-operativity of the ion permeation mechanisms within the TRPM5 channel, we used PENSA to calculate the state-specific information (SSI) shared between discrete state transitions in the occupancy distributions of both of the pore binding sites ***Thomson et al.*** (***2021***); ***Vögele et al.*** (***2022***). The methodology has been described in greater detail in our previous work ***Ives et al.*** (***2023***).

In brief, a timeseries distribution with a timestep of 20 ps for each binding site was obtained. For each frame, the ion’s atom ID number was recorded if an ion occupied the binding site in this frame (occupied state). By contrast, if the binding site was unoccupied (vacant state), an ID of -1 was recorded. We then quantified by mutual-information whether ion transitions from occupied to vacant, or vice versa, at one site were coupled to similar ion transitions at the second ion binding site. To account for statistical noise that can arise from distributions even if they are uncorrelated with one another due to small-batch effects (***McClendon et al., 2009***; ***Pethel and Hahs, 2014***), we calculated a statistical noise threshold. This threshold level was subtracted from the measured SSI values to yield the excess mutual information, or excess SSI (*exSSI*) above noise.

## Results

### Cation conductance of the TRPM5 channel in di-cationic solutions

We performed *in silico* simulations of open state *Danio rerio* TRPM5 (***Ruan et al., 2021***) embedded in a dual POPC lipid bilayer system, with a di-cationic solution of 135 mM NaCl and 135 mM CaCl_2_ in the central dense aqueous compartment, and 15 mM NaCl and 15 mM CaCl_2_ in the outer diluted aqueous compartments, respectively (Figure 1). An anti-parallel CompEL double bilayer setup (***Kutzner et al., 2011***) was used to yield a bio-mimetic transmembrane voltage of ∼ -50 mV across both embedded channels, as well as higher voltages of -130 mV, -380 mV and -610 mV to increase the number of permeation events and improve the statistics of our analyses (Figure 1). The 9:1 ion concentration gradient between the middle and the outside bulk compartment acted synergistically with the membrane voltage to drive ion permeation.

Our simulations showed a continuous flow of permeating ions, resulting in a total of 374 permeation events across all investigated simulation conditions performed with the CompEL setup. Even though the ion gradient provided an additional driving force for permeation alongside the voltages, the calculated conductances from our *in silico* electrophysiology simulations, in a range between 7 and 38 pS (Table 2), were generally in good agreement with the published conductance values of 23–25 pS from *in vitro* electrophysiology experiments on TRPM5 in NaCl based solutions (***Hofmann et al., 2003***; ***Prawitt et al., 2003***).

**Table 2.** Calculated conductances and selectivities from CompEL simulations of ion permeation in the TRPM5 channel. Mean inward conductances and standard error of the mean (SEM) were calculated from overlapping 50 ns windows from three-fold replicated 500 ns simulations of an anti-parallel double bilayer system. The number of permeation events associated with the conductance for each cation is displayed in brackets below the respective conductance values. Mean selectivity ratios of Na^+^ and Ca^2+^ permeation events and SEM were calculated from three-fold replicated 500 ns simulations.

| Conc. gradient | Voltage (mV) | Conductance (pS) |  |  |  | P<sub>Na</sub>/P<sub>Ca</sub> |
| --- | --- | --- | --- | --- | --- | --- |
|  |  | Na<sup>+</sup> | Ca<sup>2+</sup> | Cl<sup>-</sup> | Overall |  |
| 135 mM :<br>15 mM | -50 | 16 ± 3.1<br>(15) | 0 ± 0.0<br>(0) | 0 ± 0.0<br>(0) | 16 ± 3.1 | ∞ |
|  | -130 | 7 ± 1.3<br>(18) | 0 ± 0.0<br>(0) | 0 ± 0.0<br>(0) | 7 ± 1.3 | ∞ |
|  | -380 | 4 ± 0.5<br>(32) | 5 ± 0.8<br>(19) | 0 ± 0.1<br>(1) | 9 ± 1.0 | 1.9 ± 0.41 |
|  | -610 | 10 ± 0.9<br>(115) | 29 ± 2.5<br>(168) | -1 ± 0.2<br>(6) | 38 ± 3.1 | 0.8 ± 0.13 |

### Low-voltage simulations in di-cationic solutions show exclusive permeation of Na^+^ through TRPM5

At the lowest simulated voltages of ∼ -50 mV and -130 mV, we observed complete Na^+^-selectivity in mixed Ca^2+^/Na^+^ solutions, with no recorded Ca^2+^ permeation during an accumulated simulation time of 1.5 𝜇s. During the same time span, 15 (-50 mV) and 18 Na^+^ ions (-130 mV) traversed the TRPM5 pore, respectively, in accordance with its general conductance level (Table 2).

Analysis of the pore architecture of TRPM5 showed no major conformational changes throughout the course of the simulations. The TRPM5 pore possesses two main constrictions: an upper constriction formed by the sidechains of Q906 and by G905 of the three-residue selectivity filter (SF), and a lower constriction formed by the sidechains of I966 of the lower gate (Figure 1). A minor constriction can also be observed ∼13 Å above the SF, in the extracellular pore vestibule (EPV) (Figure 2). This constriction is formed by the turret loop between the pore helix (PH) and the S6 helix.

**Figure 2.**
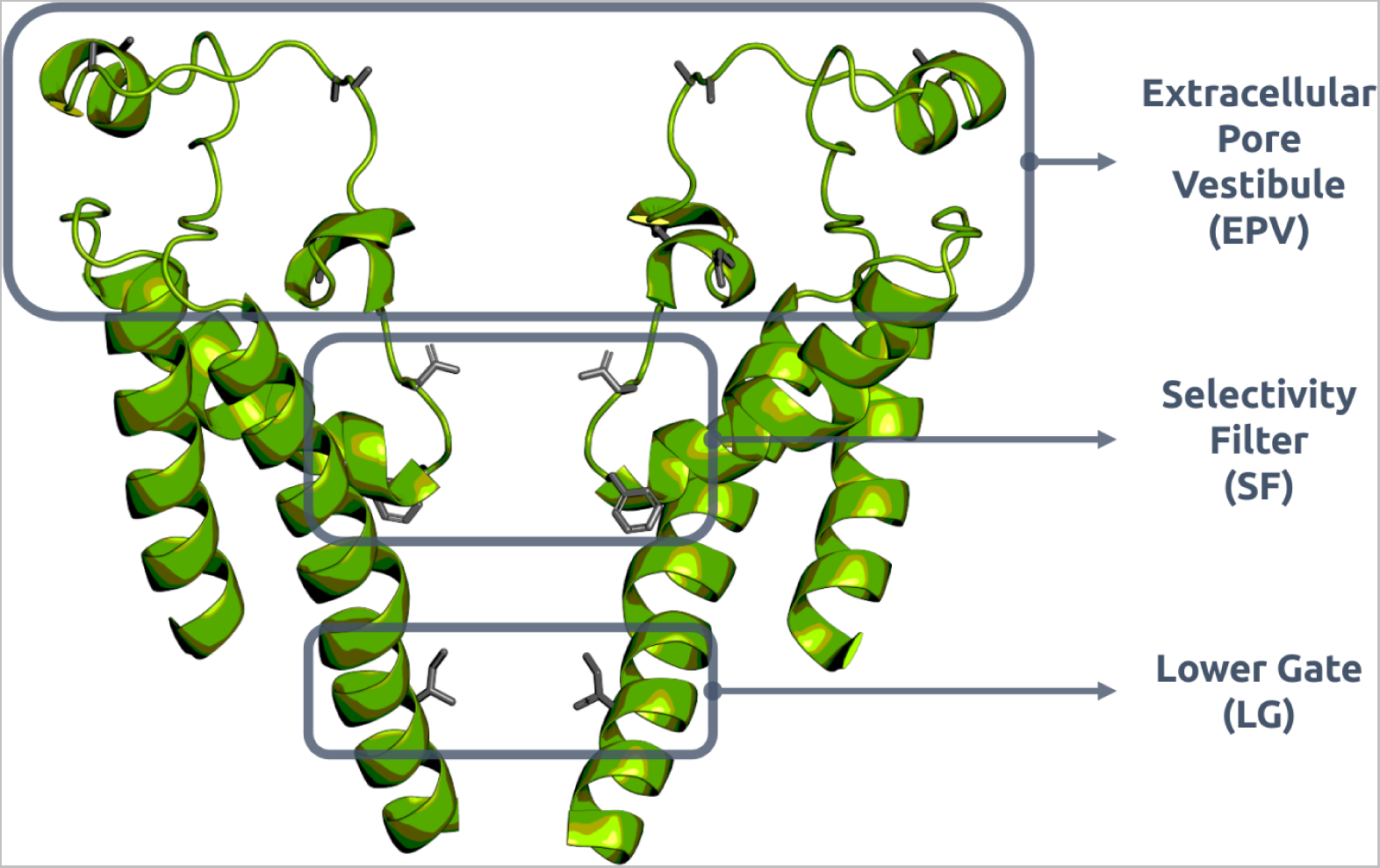
Overview of the structure of the TRPM5 channel of *Danio rerio* used in this work (two of the four subunits are omitted for clarity). TRPM5 has a short, three-residue selectivity filter (SF) consisting of Q906, G905, and F904. The hydrophobic lower gate (LG) of TRPM5 is formed by I966. In this study, the pore is defined as the region between the the two constrictions of the channel, namely Q906 of the SF and I966 of the lower gate. Above the pore is the extracellular pore vestibule, which contains a number of acidic residues, such as E910, E911, D919, D920, D925, and E928. All residues mentioned by name are displayed as grey sticks.

In our simulations at -50 mV and -130 mV, Na^+^ cations first entered the EPV region of the TRPM5 pore, where they showed a broad association with the protein matrix. Permeating Na^+^ cations then traversed the SF rapidly, and entered the pore cavity. They spent a substantial amount of time occupying the cavity before passing through the lower gate and exiting the pore at the intracellular face.

As opposed to monovalent Na^+^, Ca^2+^ ions did not readily enter the inner pore of TRPM5 during the course of the simulations. Ca^2+^ cations chiefly occupied the EPV region at the extracellular entrance (see Figure 2). 3D density maps of Na^+^ and Ca^2+^ ions further confirmed this observation (Figure 3). The maps show substantial Ca^2+^ density in the EPV, particularly near the acidic residues on the loop between the PH and the S6 helix, namely: E910, E911, D919, D920, D925, and E928. We observed that Ca^2+^ ions occasionally migrated from the EPV toward the pore, however they were blocked from entering the cavity at the SF, particularly at the constriction formed around G905 and F904 from each subunit (Figure 3).

**Figure 3.**
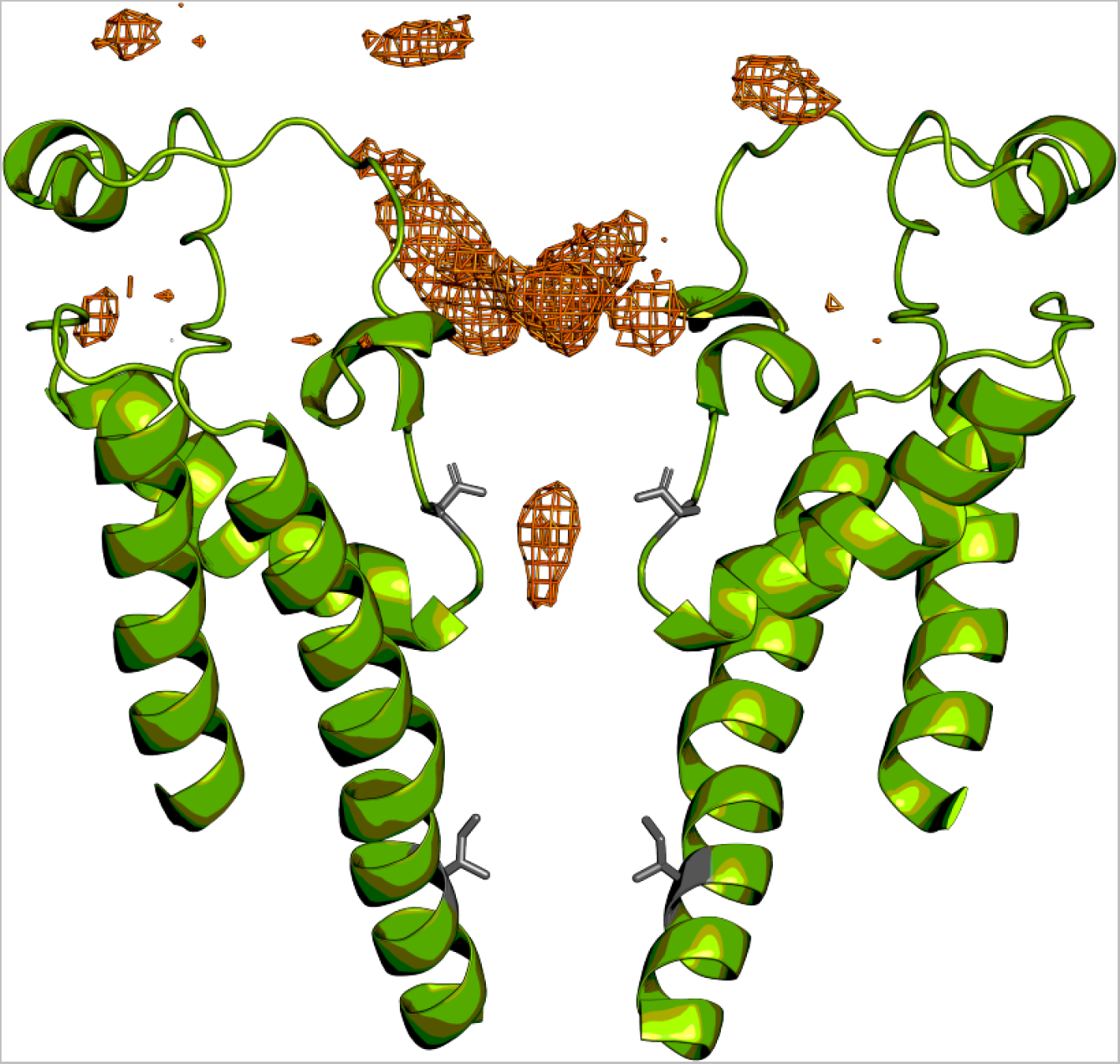
3D density map of Ca^2+^ cations around the EPV and SF of TRPM5. The density of Ca^2+^ ions was calculated from concatenated trajectories of TRPM5 in a di-cationic solution under a transmembrane voltage of ∼ -50 mV generated by the CompEL method. A major density maximum is seen within the EPV, where Ca^2+^ associates. Occasionally, Ca^2+^ ions migrated into the SF (minor density maximum at Q906); however, they were not able to traverse past the SF at bio-mimetic voltages. The sidechains of Q906 of the SF and I966 of the lower gate are shown as sticks in grey.

### Voltage dependence of simulated TRPM5 ion selectivity

As the membrane voltage was increased in our CompEL simulations, we observed the Na^+^ selectivity (P_Na_/P_Ca_) to be diminished (Table 2). At a voltage of both ∼ -50 mV and ∼ -130 mV, we recorded complete Na^+^-selectivity, with no Ca^2+^ permeation events in any of the simulations. At a voltage of ∼ -380 mV, the *in silico* electrophysiology simulations continued to display slightly Na^+^-selective permeation; however, when the voltage was further increased to ∼ -610 mV, the Na^+^-selectivity was lost. Furthermore, higher-voltage simulations also yielded a small number of Cl^-^ permeation events, with anions permeating through to the extracellular solution.

Our findings suggest relatively weak cation binding sites within the pore domain, in line with the absence of negatively charged residues lining the SF and inner cavity, due to the substantial effect of supra-physiological membrane voltages. To further explore the ion permeation dynamics in TRPM5 and their underlying determinants, we thus aimed to enhance the sampling of both Na^+^ and Ca^2+^ permeation, while at the same time remain within the Na^+^-selective voltage regime. We selected an intermediate voltage of ∼ -340 mV for investigating permeation in mono-cationic solutions to ensure a sufficient number of traversals of both Ca^2+^ and Na^+^ ions in the monovalent-selective regime.

### Mechanistic insights into ion permeation in TRPM5 from mono-cationic solutions

We conducted *in silico* electrophysiology simulations with an applied electric field, generating a membrane voltage of ∼ -340 mV, to investigate the permeation mechanism of Na^+^, K^+^ and Ca^2+^ ions in mono-cationic solutions at sufficient sampling efficiency (Table 3). As shown in Figure 4, we observed a clear difference between the behaviour of monovalent Na^+^ and K^+^ ions in the channel and the divalent Ca^2+^ ions, especially near and in the central cavity.

**Figure 4.**
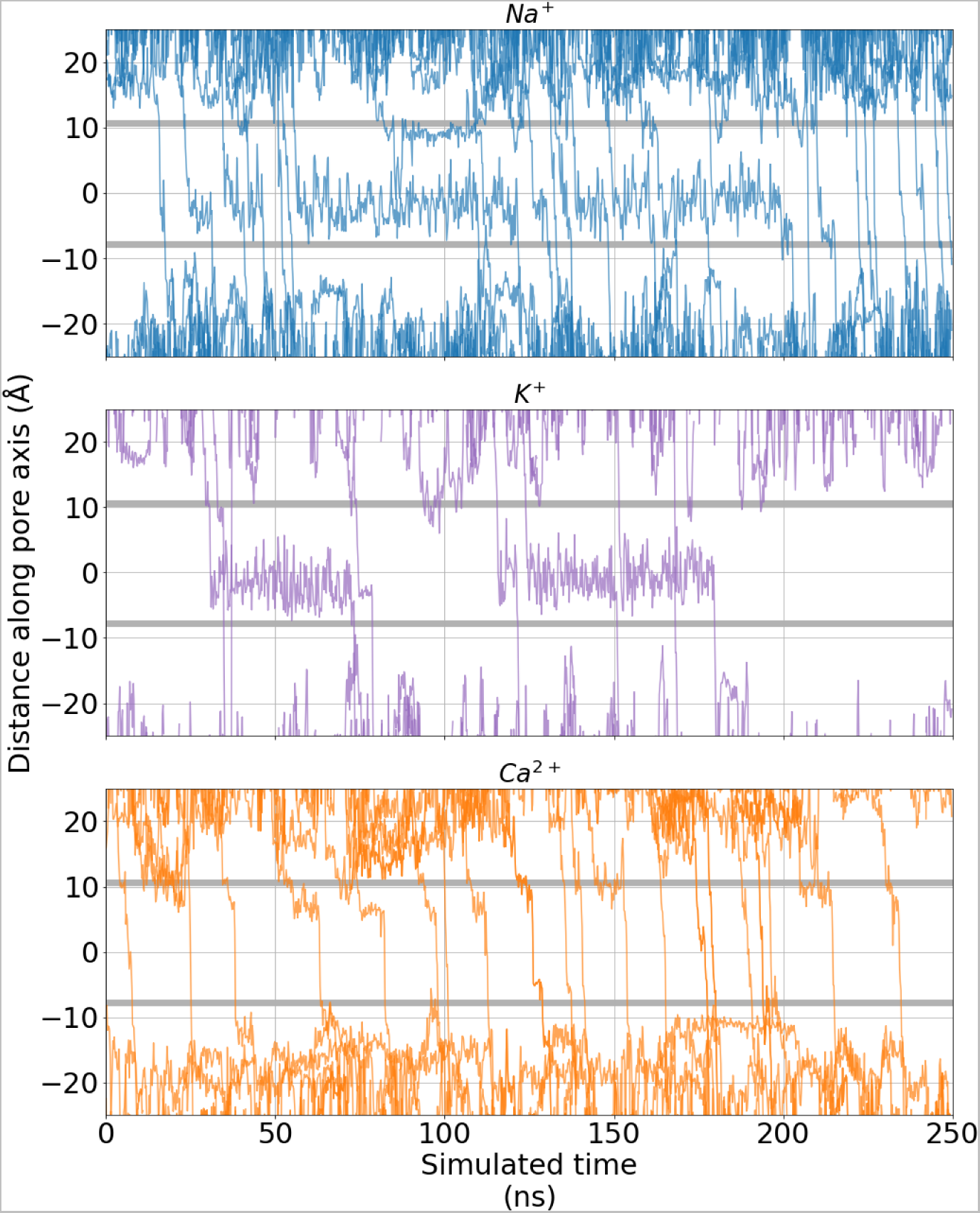
Exemplar permeation traces of the 𝑧-coordinate of permeating Na^+^ (*blue, top*), K^+^ (*purple, middle*), and Ca^2+^ (*orange, bottom*) over time, plotted from simulations performed in a mono-cationic solution with an applied electric field (-340 mV). The shaded grey regions represent the average position of the pore constrictions formed by Q906 in the SF (*upper*) and I966 of the hydrophobic gate (*lower*). Please note, only cations that fully permeate through the pore within the 250 ns simulations are shown in the plot.

**Table 3.**
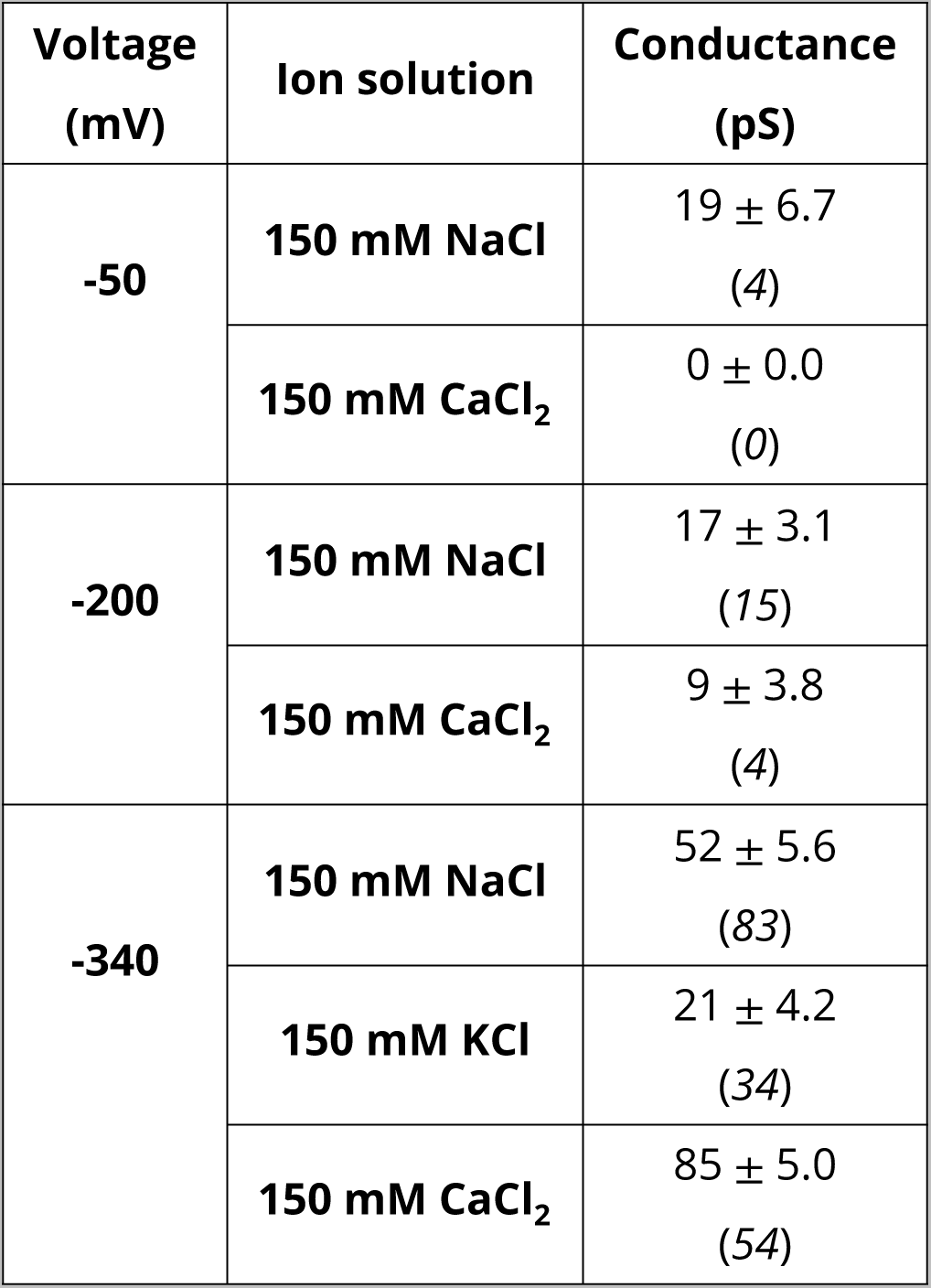
Calculated conductances from applied field simulations of ion permeation in the TRPM5 channel. Mean inward conductances and standard error of the mean (SEM) were calculated from overlapping 50 ns windows from three-fold replicated 500 ns simulations of a single bilayer system. The number of permeation events observed for each cation is displayed in brackets below the respective conductance values.

As can be seen, whereas Na^+^ and K^+^ ions occupied the central cavity of the channel for most of the simulated time, permeating Ca^2+^ ions traversed the inner cavity rapidly, not exhibiting any apparent immobilisation within the cavity. Despite occupying the cavity for extended periods of time, Na^+^ and K^+^ ions did not seem to bind to a particular binding position or residue within the cavity, but instead explored nearly the entire cavity volume before they permeated through the lower gate to the intracellular side. In this way, the cavity serves to store a monovalent ion rather than providing specific binding sites for it. A similar behaviour has recently been described for simulations of Na^+^ ions in the cavity of the homo-dimeric, endo-lysosomal Na^+^-selective cation channel TPC2 (***Milenkovic et al., 2021***).

Looking at the density of ions along the pore axis, and using the negative logarithmic density as an estimate for the underlying free energy profile at the examined non-equilibrium permeation conditions under a membrane voltage of -340 mV, it can be observed that the cavity region formed only a shallow, broad energy minimum for the permeating monovalent cations, whereas in contrast, permeating Ca^2+^ ions experienced a small apparent energy barrier in the same region (Figure 5). Both monovalent and divalent ions showed further binding to a relatively shallow binding site at the EPV. In addition, all ion types experienced a slight energy barrier to translocation near the intracellular channel exit (hydrophobic lower gate). Notably, the ions did not show major interactions with the SF. This observation, again, is in accordance with observations made in simulations of TPC2 (***Milenkovic et al., 2021***).

We conducted additional simulations using applied external electric fields of differing magnitudes. Similar to the low-voltage CompEL simulations, whereas the main features of the ion density and free energy estimates occurred across all tested voltages, Ca^2+^ was increasingly excluded from the cavity at these lower voltages, and no longer able to enter into the cavity at the lowest voltage of -50 mV during the time span of our simulations (Figure S1). This again showed that the ion selectivity of TRPM5 was voltage-dependent in the simulations. We therefore next aimed to elucidate the molecular foundations of this behaviour and, importantly, the ion selectivity of TRPM5 in general.

### Solvation profiles of cations during channel permeation

To probe if cation desolvation played a part in selective ion permeation, we calculated the number of water oxygen atoms within a 3 Å radius around the ions, representing their first solvation shell (Figure 5). Amongst other mechanisms (***Ives et al., 2023***; ***Zhang et al., 2023***), the desolvation of permeating ions has previously been reported to represent an important potential selectivity mechanism in ion channels (***Noskov and Roux, 2007***; ***Kopec et al., 2018***). Differences in the desolvation energies of permeating ions provide a thermodynamic penalty which can underpin the more favourable permeation of an ionic species over another. Here, the free energy required to desolvate Ca^2+^ strongly exceeds that for Na^+^ and K^+^ (***Marcus, 1991***), such that this difference could give rise to monovalent-selectivity in TRPM5.

**Figure 5.**
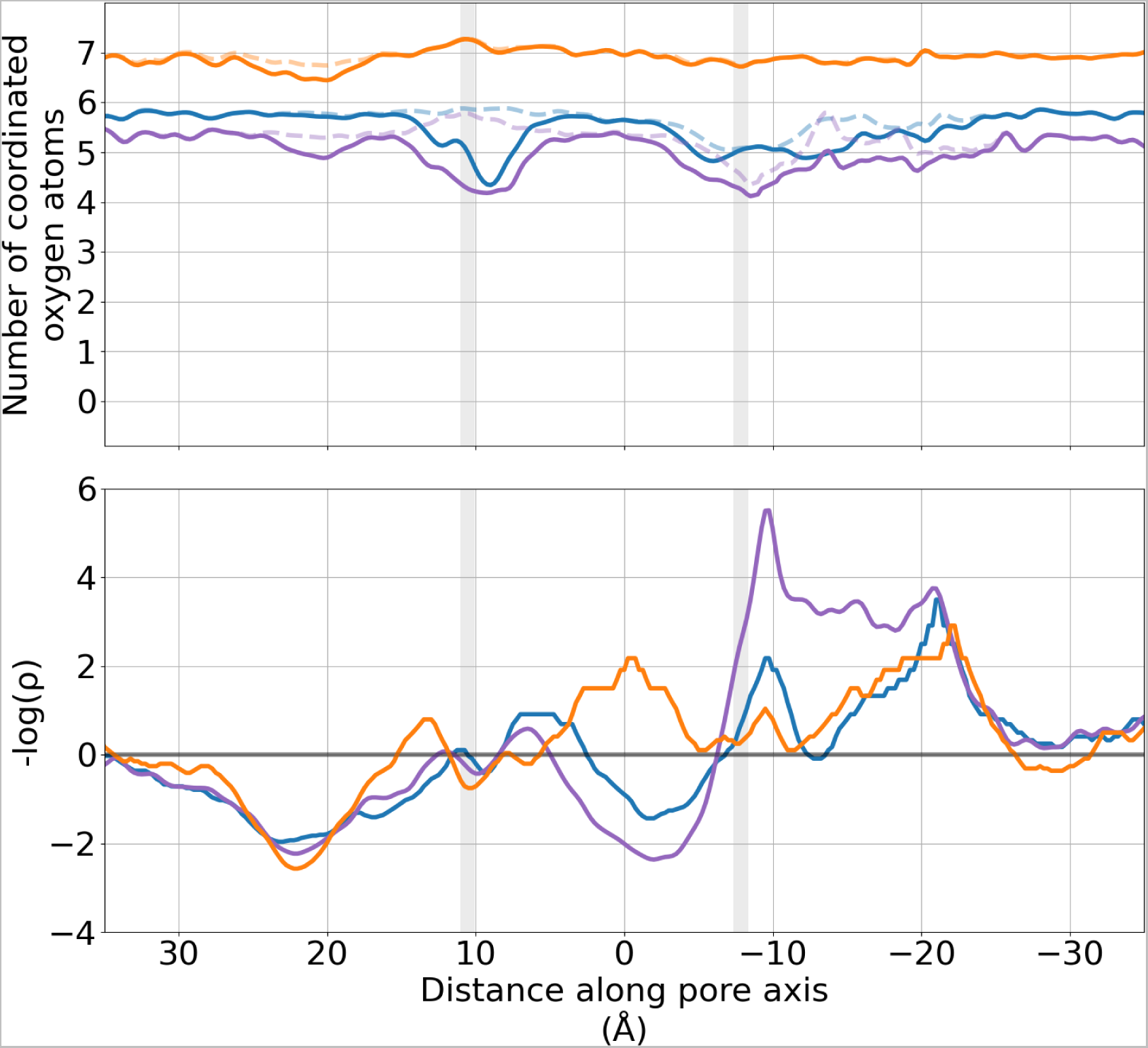
Solvation and log-density profiles of Na^+^ (*blue*), K^+^ (*purple*) and Ca^2+^ (*orange*) cations in the TRPM5 pore from simulations with a mono-cationic solution, and an applied transmembrane voltage of ∼ -340 mV. (a) The mean number of oxygen atoms of water molecules (*solid line*) and of protein oxygen atoms (*dashed line*) within 3 Å of each permeating cation is shown. (b) Negative logarithmic density profiles of permeating cations as estimates of the non-equilibrium energy surface the ions experience in the pore (energy unit: 𝑘_𝐵_ T). Minima reflect binding sites, while maxima indicate barriers between the binding sites. The location of the pore constrictions formed by Q906 (*upper*) and I966 (*lower*) are shown as grey regions. The curves in both plots have been smoothed using a Gaussian filter with a sigma value of 2.

In the bulk solution of the simulated systems, Na^+^, K^+^, and Ca^2+^ ions showed the expected water coordination number of their solvation shells. As both Na^+^ and K^+^ ions entered the pore of TRPM5, they became partially desolvated by Q906, with its side chain displacing 1–2 water molecules from the first solvation shell of the ions. After traversing the constriction at Q906, the monovalent ions were then resolvated in the pore cavity, before again being partially desolvated at the hydrophobic lower gate formed by I966. By contrast, the rapidly permeating Ca^2+^ ions did not show any significant desolvation when they crossed the SF, cavity, or hydrophobic lower gate of TRPM5.

The solvation profiles of permeating cations displayed an additional region of differing desolvation within the EPV region, highlighted previously (Figure 2). In this region, Ca^2+^ and K^+^ ions were partially desolvated, indicating closer interactions with the acidic residues in the EPV region. By contrast, Na^+^ cations did not show any desolvation in this location. We observed similar solvation profiles for permeating cations in both our simulations using an external applied electric field in mono-cationic solutions (Figure S2) and in the CompEL simulations in di-cationic solutions (Figure S3), across a range of voltage magnitudes.

Summarising, these findings suggest that ion desolvation in the SF or inner pore is not a major factor in achieving selectivity for monovalent cations. Since both monovalent and divalent cations occupied the EPV, filtering for monovalent ions must occur later in the permeation pathway. However, Ca^2+^ ions were not desolvated when they traversed the inner cavity. As the energetic penalty for desolvating Ca^2+^ is far larger than for Na^+^ or K^+^ (***Marcus, 1991***), the observed desolvation profiles therefore suggest that desolvation does not underpin the deselection of Ca^2+^ ions in TRPM5.

### Why does the central cavity form an attractive site for monovalent cations but a repulsive site for divalent cations?

The presence of a water-filled internal cavity is a conserved feature amongst cation-selective channels. The cavity serves to maintain a high degree of ion hydration despite locating to the centre of the hydrophobic lipid bilayer, and to focus the membrane voltage difference onto the SF (***Doyle et al., 1998***). Like in other cation channels, we observed in TRPM5 that the major permeating species, Na^+^ and K^+^ ions, were re-hydrated and transiently captured in the cavity, following their permeation through the SF (***Milenkovic et al., 2021***). At higher voltages, Ca^2+^ ions were able to enter and traverse the cavity, but did not alter their hydration number during this process. This suggests that they did not interact favourably with any of the cavity-lining residues or the cavity overall. At lower voltages, by contrast, Ca^2+^ ions were completely excluded from entering the cavity.

We therefore investigated the difference between the pore and cavity properties of the highly Ca^2+^-selective TRP channel, TRPV5, and the monovalent-selective TRPM5. Contrary to TRPM5, the cavity of TRPV5 shows a high occupancy with Ca^2+^ ions (***Ives et al., 2023***). As displayed in Figure 6, the general features of the pore are preserved with a constriction at the extracellular SF, a wider internal cavity region, and a second constriction at the intracellular gate. TRPM5 has a markedly shorter SF, while its cavity is wider than that of TRPV5. However, there is a substantial difference in the pore lining of the two TRP channels. Whereas the TRPV5 SF is a strongly hydrophilic region, TRPM5 does not display increased hydrophilicity within its SF. The transition from the SF to the cavity is slightly hydrophobic in TRPM5, while this is a hydrophilic region in TRPV5. There are no differences between the hydrophobicity of the two channels at the intracellular gates.

**Figure 6.**
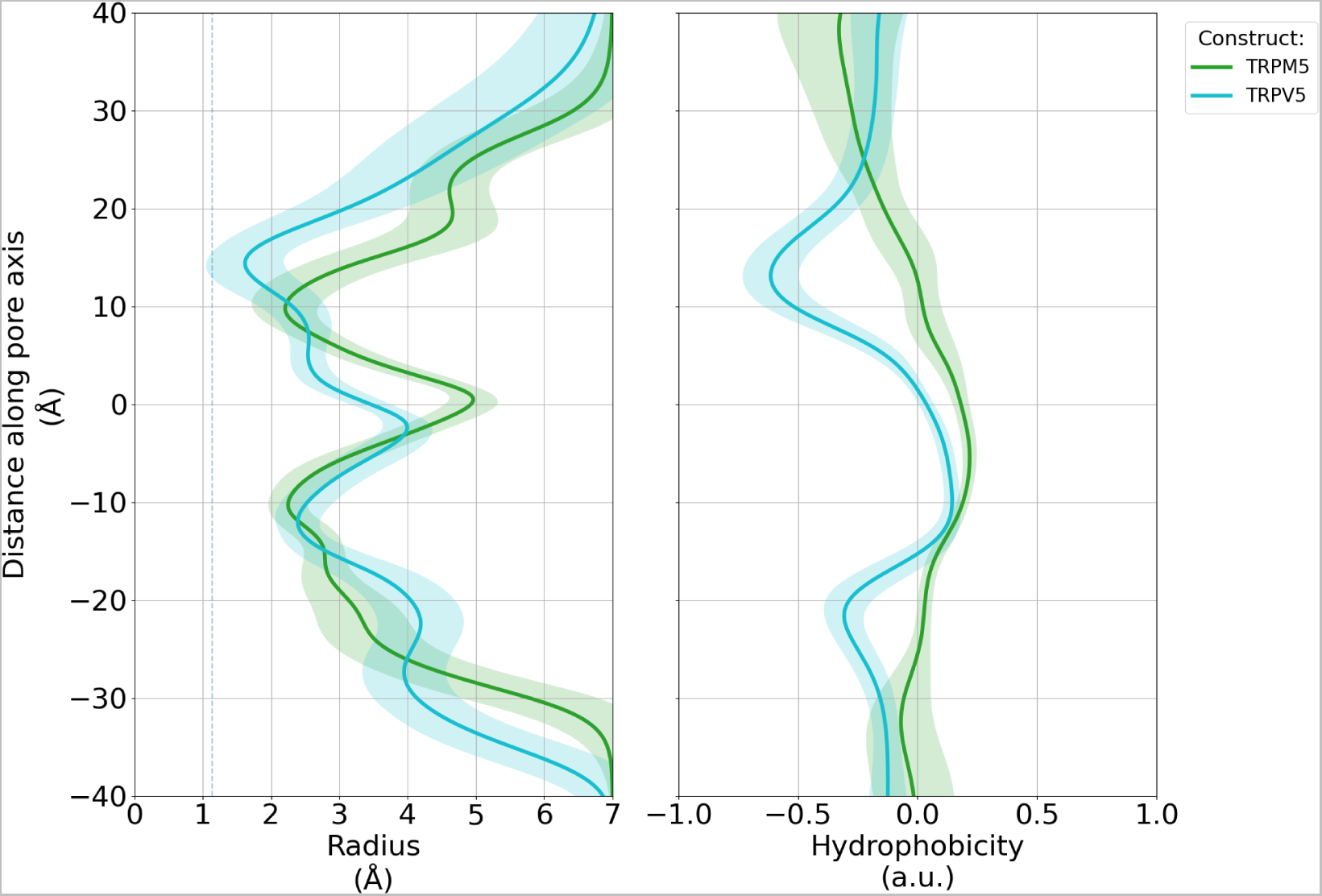
Pore architecture of the monovalent-selective TRPM5 channel (*green*) and the Ca^2+^-selective TRPV5 channel (*cyan*) from MD simulations. The average pore radius (***a***) and hydrophobic profile (***b***) for each channel was calculated using CHAP (***Rao et al., 2019***). The standard deviation is shown as shaded regions. The profile of the TRPV5 was generated from simulation data previously published (***Ives et al., 2023***). The shaded grey regions represent the average position of the pore constrictions formed by Q906 in the SF (*upper*) and I966 of the hydrophobic gate (*lower*) in TRPM5.

The differing properties of the inner pore (cavity and SF) suggest that the absence of a favourable interaction of TRPM5 with Ca^2+^ in this region arises due to the raised hydrophobicity of its SF and upper portion of its inner cavity (hydrophobic funnel). In particular, the transition zone between the SF and the cavity in TRPM5 is lined by large hydrophobic residues at the bottom of the SF, especially F904 and I903. This sequence is shared with TRPM4, which is also a monovalent-cation selective channel (Fig. 7A). On pore-forming helix S6 of TRPM5, the additional hydrophobic residues V959 and L962 line the cavity towards the hydrophobic lower gate at I966, whereas only two polar side chains, N958 and N962, are involved. The conservation level of the large hydrophobic residues lining the cavity is generally high (dark and light pink surface colour in Fig. 7B).

**Figure 7.**
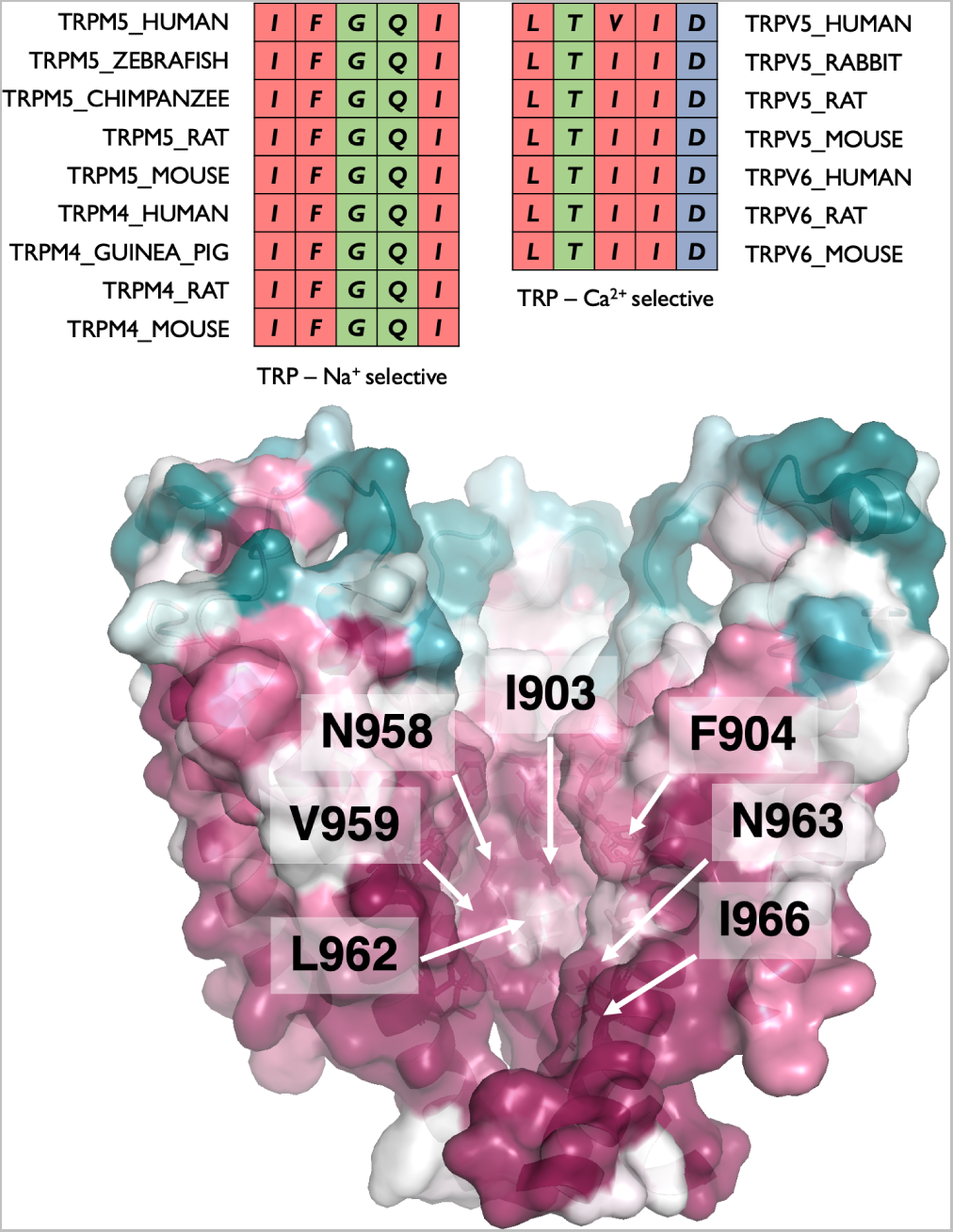
(Top) Sequence conservation of the SF and transition zone to the inner cavity in the monovalent-selective channels TRPM5 and TRPM4 compared to the Ca^2+^-selective channels TRPV5 and TRPV6. Colours according to Clustal Omega convention (***Sievers et al., 2011***). (Bottom) Evolutionary conservation of the pore cavity in TRPM5 channels. Evolutionary conservation scores were calculated using ConSurf (***Yariv et al., 2023***). The evolutionary conservation scores were projected onto the structure of TRPM5 from *D. rerio*, with one subunit removed for clarity. Figure made with Pymol (***DeLano, 2002***).

The energetic cost of placing monovalent cations into a hydrophobic environment is smaller than that for divalent cations, even when they retain their hydration shells. The Born energy for divalent cations, for example, quantifying this electrostatic energy penalty in continuum models, is four times larger than the penalty associated with monovalent cations (***Born, 1920***). As observed in the simulations under increased voltage, the protein matrix does not form favourable interactions with Ca^2+^ ions in its inner pore. The hydrophobicity of the TRPM5 pore gradually increases along the pore axis, with only few hydrophilic sites within the SF or central cavity.

Our results thus suggest that Ca^2+^ ions are unable to enter the increasingly hydrophobic environment of this region at physiological voltages, and only penetrate past the SF under high-voltage conditions, above the physiologically relevant level.

### Selectivity for monovalent cations is linked to permeation co-operativity between two binding regions

We hypothesised that, due to the presence of an additional binding or ’storage’ region for monovalent cations in the internal cavity compared to Ca^2+^, the permeation mechanism for monovalent ions may be more efficient than for Ca^2+^. In previous work, we developed a mutual-information based quantification method of the level of cooperativity when ions permeate across multiple ion channel binding sites, termed state-specific information (SSI; (***Thomson et al., 2021***; ***Vögele et al., 2022***; ***Ives et al., 2023***)). In brief, SSI quantifies the probability that a state change of one binding site, such as a change from binding an ion to becoming vacant upon ion permeation, is correlated to a similar state change in a second binding site, in which case the unbinding events are coupled to one another for instance by a knock-on mechanism.

We applied the SSI approach to ion conduction in TRPM5, focusing on the pair of ion binding regions at the EPV and the channel cavity (see Figure 2). These two binding areas are shallow and relatively distant to one another, but locate directly to the main pore axis. In addition, they show moderate to high occupancy with monovalent cations, respectively (Figure 8).

**Figure 8.**
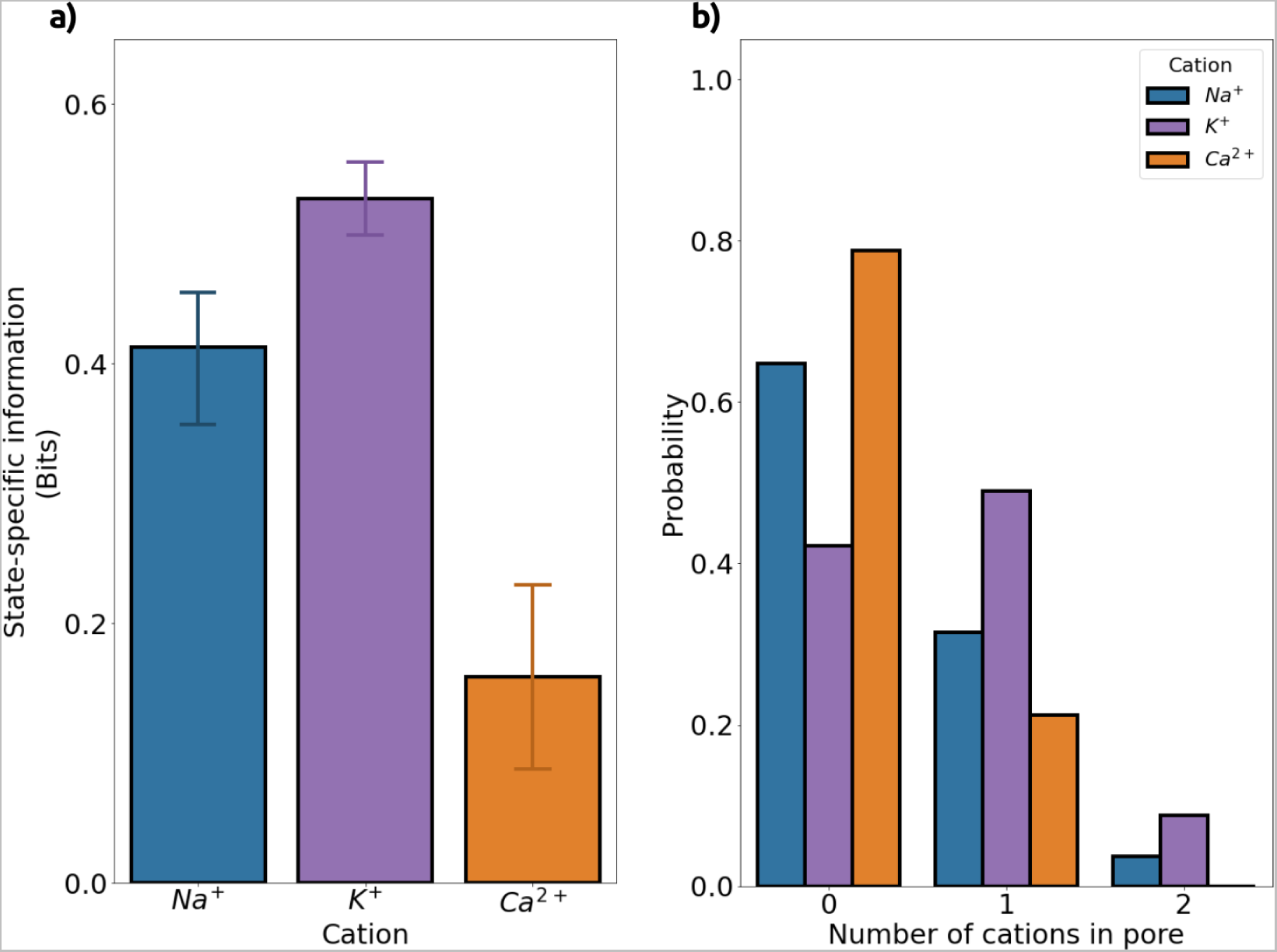
State-specific information (SSI) of cation transitions between binding sites and the average number of each cation within the pore of TRPM5. (a) Excess state-specific information (*exSSI*) between the EPV and pore cavity ion binding sites, quantifying the degree of co-operativity in the permeation mechanisms of Na^+^ (*blue*), K^+^ (*purple*), and Ca^2+^ (*orange*) ions. The mean exSSI and SEM between transitions from the two binding sites were calculated from simulations performed in mono-cationic solutions with an externally applied voltage of ∼ -340 mV. (b) Mean probability for the number of each cationic species within the inner TRPM5 pore, calculated from non-overlapping 50 ns windows from three-fold replicated 250 ns simulations. We defined the inner pore as the region between the constrictions formed by Q906 in the SF and I966 at the lower hydrophobic gate.

By using SSI, we found that both Na^+^ and K^+^ ions displayed a high level of correlation between binding and unbinding at the two successive sites, whereas the permeation of Ca^2+^ ions showed only a low degree of correlation slightly above the noise level (Figure 8). This suggests that a distant knock-on mechanism is in operation between incoming monovalent ions, which, when unbinding from the EPV bind transiently at the SF, as well as over substantial time spans within the cavity. In other words, the cavity serves as a reservoir, more likely to release a Na^+^ or K^+^ ion to the cytoplasm when a further monovalent cation approaches and inserts into the TRPM5 pore.

By contrast, Ca^2+^ ions permeated on their own. Their traversal is likely to be driven solely by the supra-physiological transmembrane electric field. Accordingly, lower-voltage simulations did not show any permeating Ca^2+^. This selectivity for monovalent ions was abolished when the driving force for Ca^2+^ permeation exceeded a certain threshold. Permeation at higher voltages could thus be described as a *’pull-through’* of Ca^2+^ ions across the otherwise unfavourable environment of the inner cavity for divalent ions. This voltage-driven *’pull-through’* occurs due to the higher charge of Ca^2+^, doubling the Coulombic driving force compared to the monovalent ions, but is unlikely to be physiological.

Our results suggest that the occurrence of two binding regions for monovalent ions, one of which is a reservoir site in the central cavity, enhance permeation efficiency via a distant knock-on mechanism. By contrast, Ca^2+^ ions are prevented from entering the pore cavity at physiologically relevant membrane voltages by a hydrophobic gate region, abolishing the reservoir binding within and thus disrupting any sizeable permeation cooperativity.

### The transition zone from the selectivity filter to the central cavity is key for monovalent cation selectivity in TRPM5

To test the hypothesis that the hydrophobicity of the transition zone connecting the SF to the central cavity provides the primary barrier to the permeation of divalent cations, we performed applied-field simulations of TRPM5 with a F904T mutation, which increases the hydrophilicity of this area, at two voltage magntitudes. In the Ca^2+^-selective TRPV channels TRPV5 and TRPV6, a thre-onine residue is located at the structurally equivalent position. Since they also contain a charged glutamate in their selectivity filter, which shows a high affinity for Ca^2+^ but is absent in TRPM5, we did not expect the TRPM5 F904T mutant to show a similarly high degree of Ca^2+^-selectivity. Rather, we hypothesised that the added hydrophilicity of the mutant may reduce the height of the hydrophobic barrier for both monovalent and divalent cations, that is, increase the flow of Na^+^ while allowing the passage of Ca^2+^ in physiologically relevant voltage ranges.

Three-fold replicated simulations of the F904T mutant in 150 mM CaCl_2_ at -130 mV showed 13 completed Ca^2+^ permeation events within a total time of 750 ns. In the same time span, 27 Na^+^ ions traversed the mutant channel in 150 mM NaCl solution (Table S3). The permeation numbers correspond to a selectivity of P_Na_/P_Ca_ of ∼2, otherwise not observed below a voltage of -380 mV. Additional control simulations at -200 mV in mono-cationic solution displayed 15 Na^+^ and 4 Ca^2+^ permeation events in the WT (Table S2), whereas the mutant conducted 88 Na^+^ ions and 28 Ca^2+^ ions within the same accumulated time span (Table S3).

These results show that the F904T mutation indeed facilitated the permeation of Ca^2+^, while at the same time increasing the Na^+^ flux. We conclude that the hydrophobic transition zone between the SF and the central cavity plays the major role in governing the selectivity of TRPM5 for monovalent cations at physiological voltages.

## Discussion

The TRP channel superfamily encompasses a broad range of cation-selective ion channels of great physiological and biomedical importance (***Ramsey et al., 2006***; ***Khalil et al., 2018***). While most members of the superfamily translocate both monovalent cations and divalent Ca^2+^ at similar permeabilities, TRPV5 and TRPV6 are strongly Ca^2+^-selective (***Ives et al., 2023***) and TRPM4 and TRPM5 are selective for monovalent cations (***Owsianik et al., 2006***). In contrast to most other Na^+^-selective channels (***Dudev and Lim, 2014***), the SFs of TRPM4, TRPM5, and the endo-lysosomal TPC2 do not contain charged residues, while they retain a relatively high abundance of hydrophobic residues (***Ruan et al., 2021***; ***Milenkovic et al., 2021***).

Ion selectivity is usually linked to the presence of specific binding sites in the channels’ SF and inner pore (***Hille, 2001***; ***Zhou et al., 2001***), cooperativity between the permeation kinetics of multiple such binding sites (***Hille, 2001***; ***Köpfer et al., 2014***; ***Ives et al., 2023***; ***Derebe et al., 2011***), ion desolvation (***Noskov and Roux, 2007***; ***Kopec et al., 2018***), or size exclusion effects (***Hille, 2001***; ***Dudev and Lim, 2014***). Our simulations showed, however, that no high-affinity binding sites for cations exist within the pore of TRPM5, that only minor ion desolvation effects occur, and that size exclusion does not play a role in selectivity. The channel can conduct both monovalent and divalent cations at slightly increased membrane voltages. We observed two shallow, broad ion binding regions for monovalent cations in the TRPM5 channel; one within the EPV above the SF, and a second within the central cavity, whereas divalent cations did not interact favourably within the cavity.

Our findings suggest a new mechanism of monovalent cation selectivity, in which the combination of an uncharged, relatively hydrophobic SF and the presence of large hydrophobic side chains at the entrance to the central channel cavity determine the monovalent-selectivity of TRPM5. Since the traversal of a divalent cation through this hydrophobic funnel incurs a larger energy penalty as compared to a monovalent cation, this hydrophobic region creates a higher energy barrier for the permeation of divalent cations (***Born, 1920***). The hydrophobic energy barrier difference generated in this way is of moderate magnitude, and therefore can be overcome by divalent cations in the supra-physiological voltage range.

Under physiologically relevant voltages, this barrier and the generally largely hydrophobic character of the central cavity prevent divalent cations from entering and residing within the central cavity, whereas monovalent cations readily enter and occupy the cavity for substantial time spans. Numerous closed-state structures of TRPM4, a close monovalent-selective homolog of TRPM5, have been published within the PDB. Several of these structures include Na^+^ cations modelled within the pore cavity (***Guo et al., 2017***; ***Duan et al., 2018***). Consequently, these structures (PDB IDs 6BCJ, 6BCL, and 6BWI) suggest that the presence of a monovalent cation binding region in the inner cavity is a conserved feature amongst the monovalent-selective TRPM4 and TRPM5 channels.

Due to this additional binding region, a distant knock-on mechanism is established between an incoming monovalent cation and the monovalent cation stored within the cavity, which greatly increases permeation efficiency. By contrast, under supra-physiological voltage conditions, divalent cations are simply ’pulled through’ the hydrophobic cavity on their own, exhibiting no interactions with the cavity matrix or other cations. Our application of the mutual-information based SSI method (***Ives et al., 2023***; ***Vögele et al., 2021***) to ions permeating through TRPM5 showed only negligible cooperativity for Ca^2+^ permeation.

Finally, we examined whether a hydrophilic mutation at the entrance to the central cavity facilitated the flux of divalent cations through this region. According to the findings discussed above, this was indeed the case in the F904T mutant of TRPM5, with substantial Ca^2+^ permeation in a voltage range that did not allow for Ca^2+^ permeation in the WT channel. In line with a hydrophobic barrier that exists for both monovalent and divalent cations, but is larger for divalent cations, the flow of Na^+^ also increased in the mutant, raising its overall conductance level.

## Acknowledgments

We thank the University of Dundee I.T. services for maintenance of the School of Life Sciences high-performance computing (HPC) cluster which was utilised in this research.

## Author contributions

CMI and UZ conceived the idea and designed the computational study, CMI conducted and analysed the WT simulation data, ATŞ conducted and analysed the F904T mutant simulations, NJT analysed the SSI data, UZ supervised the work, CMI and UZ wrote the manuscript with contributions from NJT, and all authors edited and reviewed the manuscript.

## Supplementary

### Summary of MD simulations used within this study

**Table S1.** Summary of CompEL simulation details of the TRPM5 channel. All simulations were conducted in a di-cationic solution of 75 mM NaCl and 75 mM CaCl_2_. In all simulations, the Ca^2+^ cations occupying the Ca_TMD_ were modelled, and remained bound for the duration of the simulations.

|  |  |  |  |  |
| --- | --- | --- | --- | --- |
| **Protein** | TRPM5 |  |  |  |
| **Structure** | 7MBS<br>(698-1020) |  |  |  |
| **Force field** | CHARMM36m |  |  |  |
| **Water** | TIP3P |  |  |  |
| ***In silico electrophysiology methodology*** | CompEL<br>(anti-parallel with a 9:1 concentration gradient) |  |  |  |
| **Ion** | 75 mM NaCl + 75mM CaCl<sub>2</sub><br>266 Na<sup>+</sup> (CHARMM36m)<br>274 Ca<sup>2+</sup> (Zhang et al.)<br>814 Cl<sup>-</sup> (CHARMM36m) |  |  |  |
| **Independent simulations** | 3 | 3 | 3 | 3 |
| **Total simulation time (μs)** | 1.5 | 1.5 | 1.5 | 1.5 |
| **Total aggregated simulation time (μs)** | 3 | 3 | 3 | 3 |
| **Ionic ratios between compartments**<br>**(Extracellular : Intracellular)** | 239 : 27 Na<sup>+</sup><br>239 : 35 Ca<sup>2+</sup><br>681 : 133 Cl<sup>-</sup> | 239 : 27 Na<sup>+</sup><br>239 : 35 Ca<sup>2+</sup><br>680 : 134 Cl<sup>-</sup> | 239 : 27 Na<sup>+</sup><br>239 : 35 Ca<sup>2+</sup><br>677 : 137 Cl<sup>-</sup> | 239 : 27 Na<sup>+</sup><br>239 : 35 Ca<sup>2+</sup><br>673 : 141 Cl<sup>-</sup> |
| **Estimated voltage (mV)** | -50 | -130 | -380 | -610 |
| **Permeation events** | 15 Na<sup>+</sup><br>0 Ca<sup>2+</sup><br>0 Cl<sup>-</sup> | 18 Na<sup>+</sup><br>0 Ca<sup>2+</sup><br>0 Cl<sup>-</sup> | 32 Na<sup>+</sup><br>19 Ca<sup>2+</sup><br>1 Cl<sup>-</sup> | 115 Na<sup>+</sup><br>168 Ca<sup>2+</sup><br>6 Cl<sup>-</sup> |
| **Total number of permeation events** | 15 | 18 | 52 | 289 |

**Table S2.**
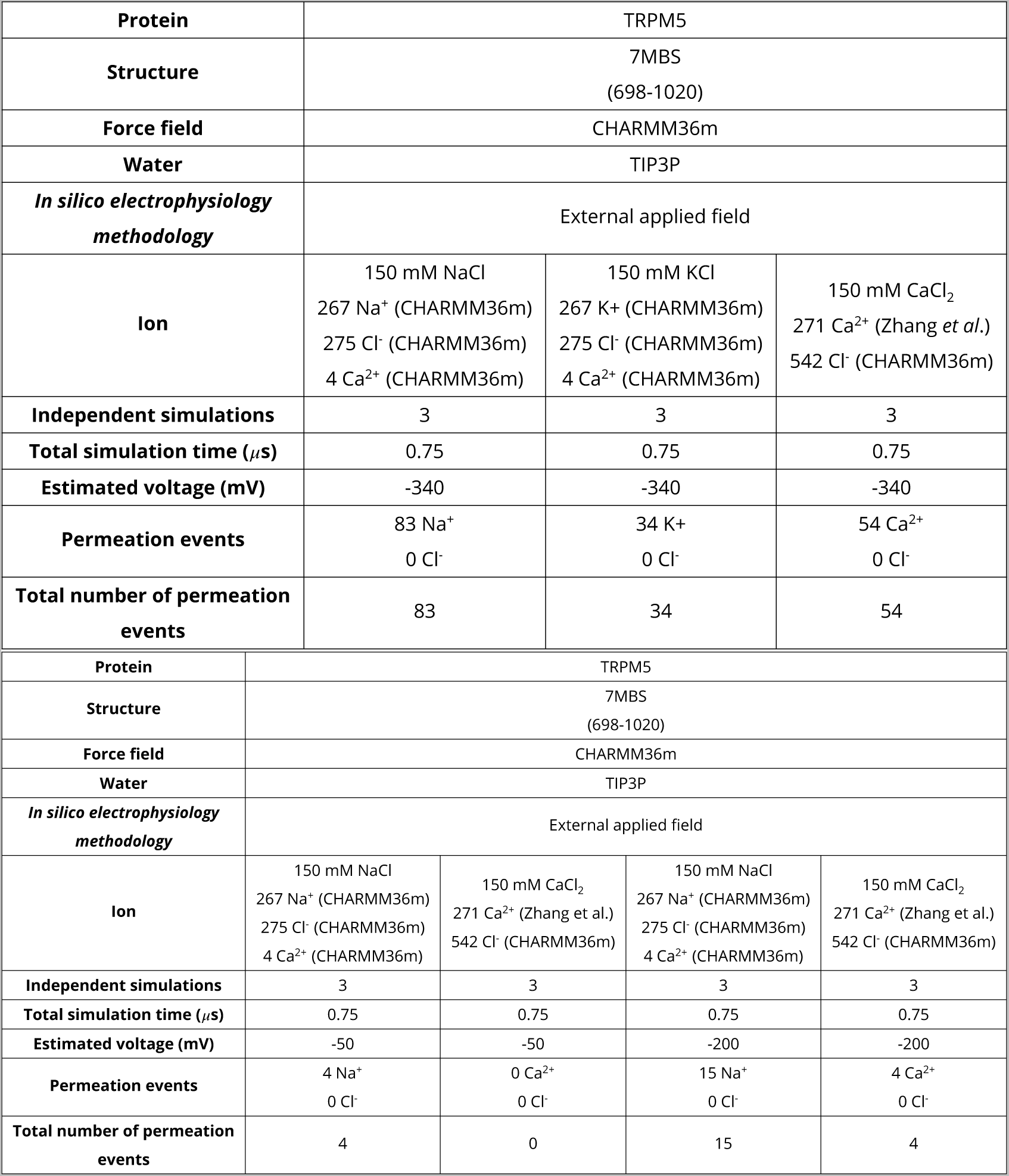
Summary of external applied field simulation details of the TRPM5 channel. All simulations were conducted in a mono-cationic solution of either 150 mM NaCl, 150 mM KCl, or 150 mM CaCl_2_. In all simulations, the Ca^2+^ cations occupying the Ca_TMD_ were modelled, and remained bound for the duration of the simulations.

|  |  |  |  |
| --- | --- | --- | --- |
| **Protein** | TRPM5 |  |  |
| **Structure** | 7MBS<br>(698-1020) |  |  |
| **Force field** | CHARMM36m |  |  |
| **Water** | TIP3P |  |  |
| ***In silico electrophysiology methodology*** | External applied field |  |  |
| **Ion** | 150 mM NaCl<br>267 Na<sup>+</sup> (CHARMM36m)<br>275 Cl<sup>-</sup> (CHARMM36m)<br>4 Ca<sup>2+</sup> (CHARMM36m) | 150 mM KCl<br>267 K<sup>+</sup> (CHARMM36m)<br>275 Cl<sup>-</sup> (CHARMM36m)<br>4 Ca<sup>2+</sup> (CHARMM36m) | 150 mM CaCl<sub>2</sub><br>271 Ca<sup>2+</sup> (Zhang *et al.*)<br>542 Cl<sup>-</sup> (CHARMM36m) |
| **Independent simulations** | 3 | 3 | 3 |
| **Total simulation time (μs)** | 0.75 | 0.75 | 0.75 |
| **Estimated voltage (mV)** | -340 | -340 | -340 |
| **Permeation events** | 83 Na<sup>+</sup><br>0 Cl<sup>-</sup> | 34 K<sup>+</sup><br>0 Cl<sup>-</sup> | 54 Ca<sup>2+</sup><br>0 Cl<sup>-</sup> |
| **Total number of permeation events** | 83 | 34 | 54 |

|  |  |  |  |  |
| --- | --- | --- | --- | --- |
| **Protein** | TRPM5 |  |  |  |
| **Structure** | 7MBS<br>(698-1020) |  |  |  |
| **Force field** | CHARMM36m |  |  |  |
| **Water** | TIP3P |  |  |  |
| ***In silico electrophysiology methodology*** | External applied field |  |  |  |
| **Ion** | 150 mM NaCl<br>267 Na<sup>+</sup> (CHARMM36m)<br>275 Cl<sup>-</sup> (CHARMM36m)<br>4 Ca<sup>2+</sup> (CHARMM36m) | 150 mM CaCl<sub>2</sub><br>271 Ca<sup>2+</sup> (Zhang *et al.*)<br>542 Cl<sup>-</sup> (CHARMM36m) | 150 mM NaCl<br>267 Na<sup>+</sup> (CHARMM36m)<br>275 Cl<sup>-</sup> (CHARMM36m)<br>4 Ca<sup>2+</sup> (CHARMM36m) | 150 mM CaCl<sub>2</sub><br>271 Ca<sup>2+</sup> (Zhang *et al.*)<br>542 Cl<sup>-</sup> (CHARMM36m) |
| **Independent simulations** | 3 | 3 | 3 | 3 |
| **Total simulation time (μs)** | 0.75 | 0.75 | 0.75 | 0.75 |
| **Estimated voltage (mV)** | -50 | -50 | -200 | -200 |
| **Permeation events** | 4 Na<sup>+</sup><br>0 Cl<sup>-</sup> | 0 Ca<sup>2+</sup><br>0 Cl<sup>-</sup> | 15 Na<sup>+</sup><br>0 Cl<sup>-</sup> | 4 Ca<sup>2+</sup><br>0 Cl<sup>-</sup> |
| **Total number of permeation events** | 4 | 0 | 15 | 4 |

**Table S3.**
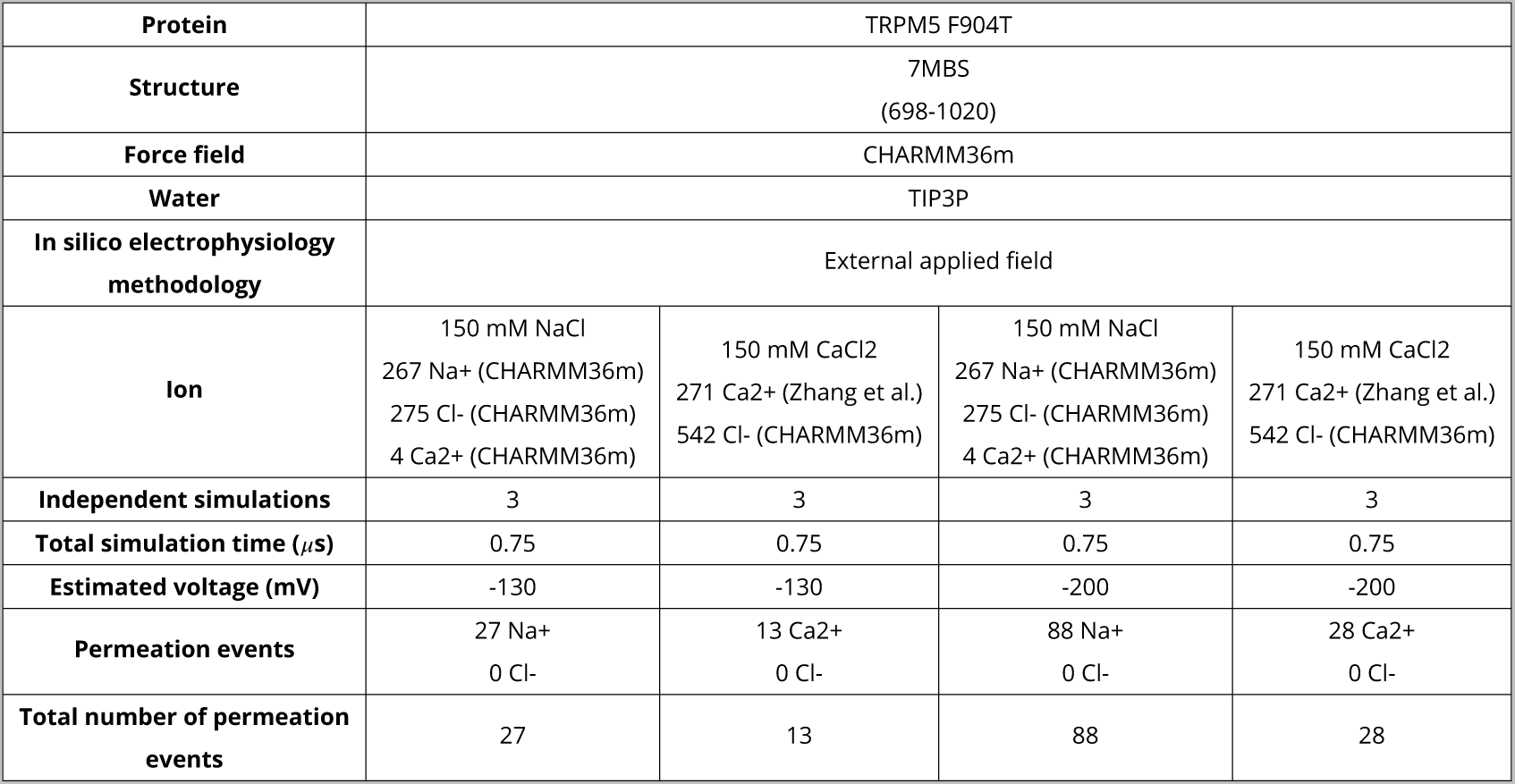
Summary of external applied field simulation details of the TRPM5 F904T channel. All simulations were conducted in a mono-cationic solution of either 150 mM NaCl, or 150 mM CaCl_2_. In all simulations, the Ca^2+^ cations occupying the Ca_TMD_ were modelled, and remained bound for the duration of the simulations.

**Figure S1.**
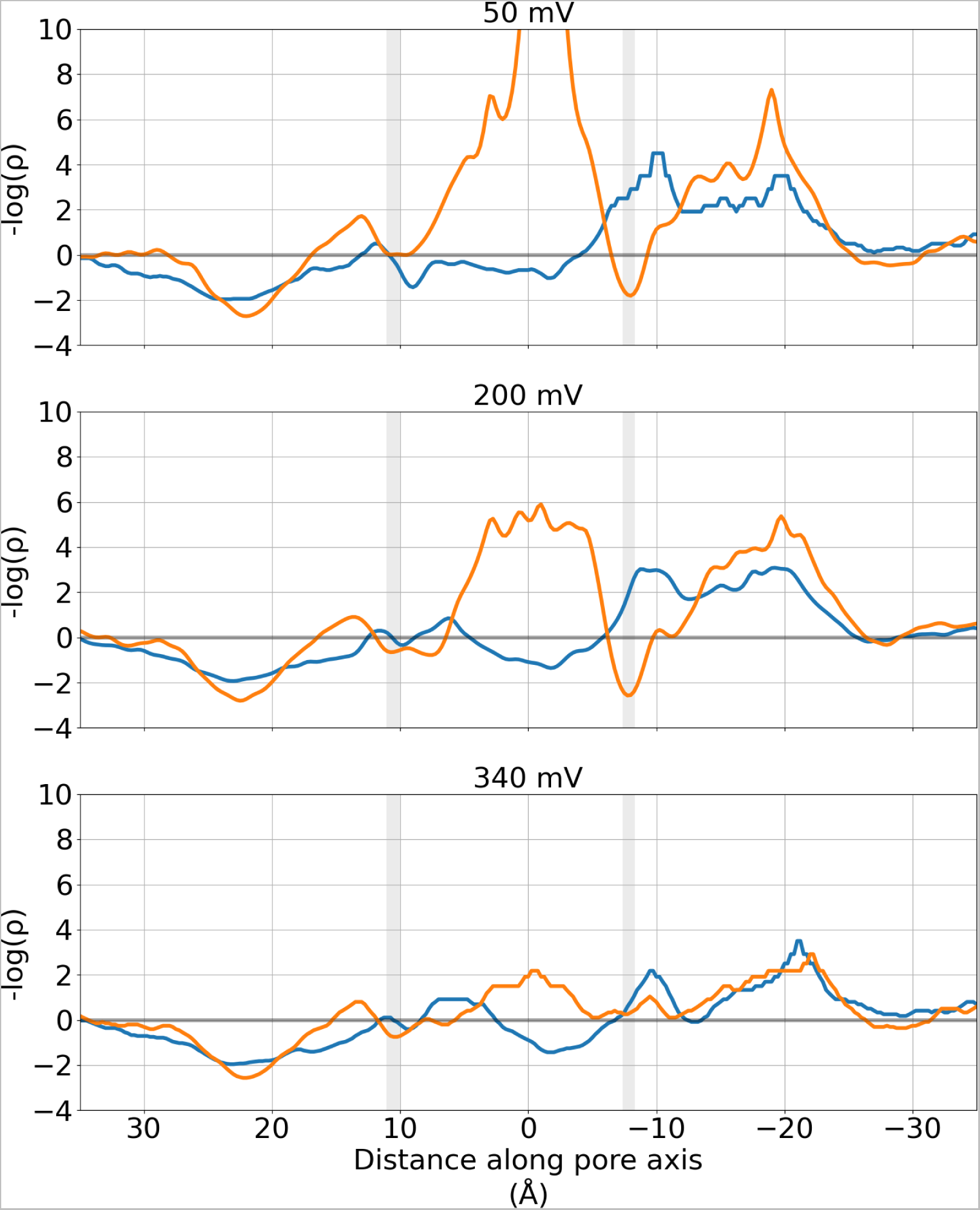
Negative logarithmic density profiles of permeating cations along the pore of the TRPM5 at different voltages. These simulations were performed in a mono-cationic solution, with an external applied electric field used to produce transmembrane voltages of ∼ -50 mV (*top*), ∼ -200 mV (*centre*), and ∼ -340 mV (*bottom*). The logarithmic ion densities represent quasi-free energies (with a nominal unit of kT). The location of the pore constrictions formed by Q906 (*upper*) and I966 (*lower*) are represented as grey regions. Both plots have been smoothed using a Gaussian filter with a sigma value of 2.

**Figure S2.**
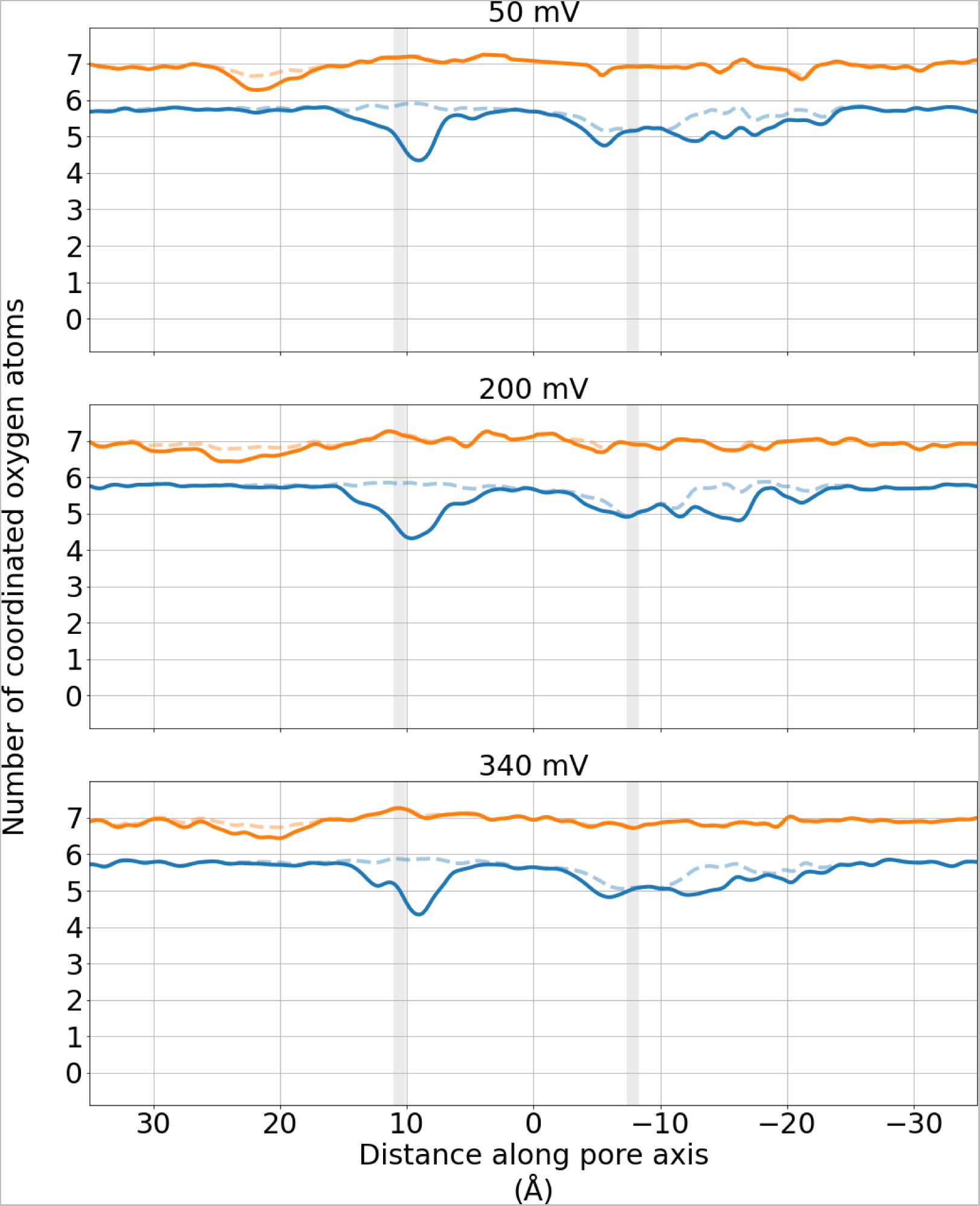
Solvation profiles of Na^+^ (*blue*) and Ca^2+^ (*orange*) cations through the TRPM5 pore. These simulations were performed in a mono-cationic solution, with an external applied electric field used to produce transmembrane voltages of ∼ -50 mV (*top*), ∼ -200 mV (*centre*), and ∼ -340 mV (*bottom*). The mean number of oxygen atoms of water molecules (*solid line*) and of any oxygen atoms of any molecule (*dashed line*) within 3 Å of each permeating cation is plotted. The location of the pore constrictions formed by Q906 (*upper*) and I966 (*lower*) are represented as grey regions. All plots have been smoothed using a Gaussian filter with a sigma value of 2

**Figure S3.**
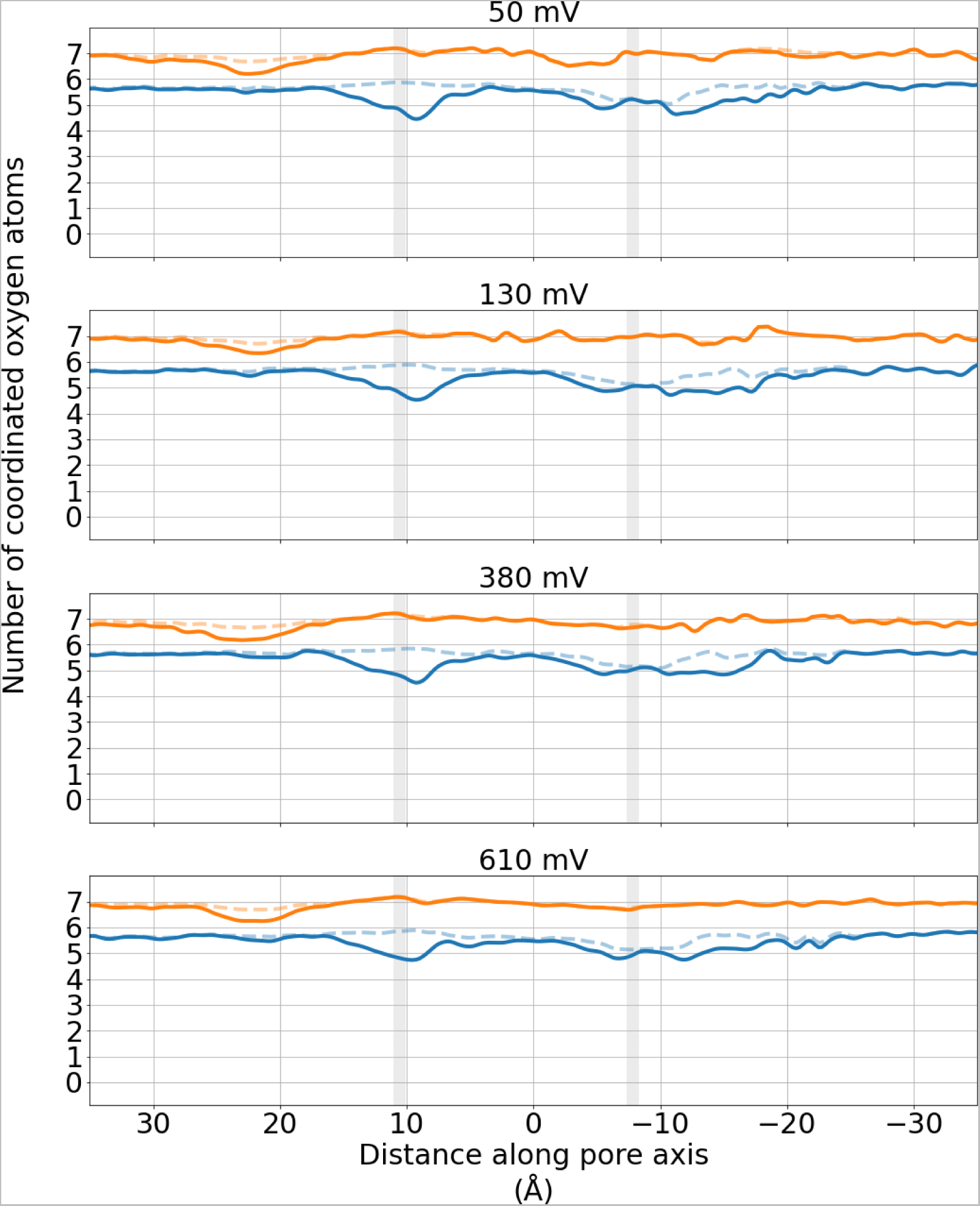
Solvation profiles of Na^+^ (*blue*) and Ca^2+^ (*orange*) cations through the TRPM5 pore. These simulations were performed in a di-cationic solution, with the CompEL methodology used to produce transmembrane voltages of ∼ -50 mV (*top*), ∼ -130 mV (*second from top*), ∼ -380 mV (*second from bottom*), and ∼ -610 mV (*bottom*). The mean number of oxygen atoms of water molecules (*solid line*) and of any oxygen atoms of any molecule (*dashed line*) within 3 Å of each permeating cation is plotted. The location of the pore constrictions formed by Q906 (*upper*) and I966 (*lower*) are represented as grey regions. All plots have been smoothed using a Gaussian filter with a sigma value of 2.

## Notes

### Competing Interest Statement

The authors have declared no competing interest.

### Summary of Updates

Manuscript updated to include additional simulations and analysis, including a TRPM5 mutant.

